# Adeno-associated Virus (AAV) Synthetic Inverted Terminal Repeat Enhances Tissue Specific Transduction and Altered Vector Induced Stress Response

**DOI:** 10.64898/2026.07.10.737493

**Authors:** Tomoko Hasegawa, Naveen Vridhachalam, Ethan Sud Nikolai, Thanu Kalikiri, Mark Ross, Robert Toennisson, Priscila Villanueva, Alicia M. Chandler, Liujiang Song, Jacquelyn J. Bower, R. Jude Samulski, Matthew L. Hirsch

## Abstract

While adeno-associated virus (AAV) vectors have shown therapeutic benefit in clinical applications, noted challenges include low transduction efficiencies, poor cellular targeting, and vector related adverse events. Recently, it was demonstrated that a rationally designed synthetic inverted terminal repeat (SynITR) altered the AAV vector-induced DNA damage response and abrogated apoptosis in human embryonic stem cells. To explore the utility of AAV-SynITR for diverse gene therapy applications, vector production, transduction, and the cellular response were evaluated in various contexts. Regarding production, SynITR preparations exhibited comparable titers to wtITR in a serotype/transgene-independent manner. Despite slightly decreased transduction efficiency in various cell lines, intravenous administration of AAV8 vectors showed SynITR enhanced transduction in a tissue-specific manner in liver (>7-fold) and kidney and pancreas (>2-fold) at equivalent vector copy numbers; however, no differences were observed in muscle/heart/spleen tissues. Interestingly, persistent γH2AX, a marker of aging/chronic inflammation, was abundant in the liver and spleen following wtITR (but not SynITR) transduction. In human corneas, SynITR enhanced transduction up to 16-fold over wtITRs. These data demonstrate that SynITRs elicit tissue-specific transduction enhancement and alter the cellular stress response. Importantly, the SynITRs offer an alternative context to elucidate wtITR biology for targeted, enhanced, and potentially safer human gene therapy.

## Introduction

Adeno-associated virus (AAV) vectors are promising tools for gene therapy and clinical data obtained from diverse applications in various tissues support the general safety of recombinant AAV (rAAV) as a gene delivery vector for genetic and acquired diseases. However, rAAV-related cellular toxicities have been reported, particularly in human stem cells, the central nervous system, and in the liver, especially when high doses are administered systemically.^1–3^ Additionally, off-target transgene expression, production costs for high titer preparations, and relatively poor transduction efficiencies limit AAV vector gene therapy applications. While capsid engineering has offered the potential to increase transduction efficiency and/or organ specificity to reduce safety concerns, many capsids fail during inter-species translational studies, which has tempered enthusiasm for relying solely on the capsid to address the limitations for AAV gene therapy.^4–6^

In addition to studies focused on more efficient production methodologies and enhanced transduction or tissue restriction via novel promoters, the AAV inverted terminal repeats (ITRs) represent a relatively understudied target for vector optimization. Following full/partial rAAV capsid uncoating, the ITRs stimulate transgenic genome processing via host repair pathways that initiate second strand synthesis and vector genome circularization thought necessary for long-term transgene expression.^7^ In fact, the ITRs are the only persistent “viral” element of rAAV gene therapy, and to date all clinical applications and FDA approved drugs rely on the natural nucleotide sequence of AAV serotype 2 [wild type ITR (wtITR)] published in 1983.^8^ Regarding rAAV production, the ITRs serve as the transgenic DNA replication origin and are implicated in vector genome packaging of the AAV capsid.^9^ However, AAV ITR biology is poorly understood and despite over 40yrs of AAV research, there remain relatively few reports of modified or novel ITRs.^10–16^ This deficit is due, in part, to technical difficulties in generating inverted DNA repeats due to their intracellular instability in *Escherichia coli*.^17–20^ Perhaps more importantly, mutations in the ITR often eliminate functions necessary for rAAV production including vector genome replication, capsid packaging, and high production titers. However, ITR mutations can also affect other physiological aspects following cellular entry such as second-strand synthesis, the durability of episomal persistence, and/or transcriptional regulation/repression/silencing, all of which could alter the transduction efficiency.^12,21^

Recently, a report highlighted the ITR’s influence on several aspects of AAV vectorology including production titers, the vector-induced DNA damage response, and the overall transduction efficiency. In that work, a rationally designed synthetic ITR (SynITR) was introduced that maintains discrete functional ITR regions, including the Rep Binding Element (RBE), the nicking stem, and the RBE’ deemed critical for vector genome replication and high titer rAAV production.^10^ The SynITR contains 25 nucleotide substitutions within the putative p53 binding sites of wtITR B and C elements that uncoupled AAV vector transduction from p53-dependent apoptosis in human embryonic stem cells (hESCs).^10^ While AAV-SynITR transduction was not statistically different from wtITR vectors in various cell lines, SynITR reporter activity was repressed >10-fold in hESCs despite equivalent capsid uncoating ^10^. Furthermore, SynITRs decreased the vector induced DNA damage response including p53 activation, demonstrating an indirect relationship between DNA damage signaling and the efficiency of transduction in hESCs.

In this work, SynITR AAV vectors were characterized in comparison to wtITR vectors for aspects relevant to the clinical application of AAV therapeutics. Importantly, SynITR AAV vector production using different capsids and transgenes were produced at similar titers of comparable quality as traditional wtITR vectors. Experiments in cultured cells of various tissue origins noted that the efficiency of SynITR vector transduction was modestly decreased for reporter function (GFP or luciferase). In stark contrast, following AAV8 intravenous injection in wt mice, SynITRs significantly enhanced transduction in the liver, kidney, and pancreas, but not in the heart, muscle, or spleen, at equivalent vector genome copy numbers. SynITR vector transduction was initially similar to wtITRs at day 3 post-injection, which continued to increase to a 7-fold elevation in the liver by the experimental endpoint (Day 23). At this experimental endpoint, evaluation of the vector-induced DNA damage response in liver tissue failed to detect canonical p53 activation or transactivation; however, persistent gamma-H2AX (γH2AX) accumulation was noted in the liver and spleen in wtITR, but not SynITR, vector treated subjects. Towards the translational relevance of SynITR vectors, transduction efficiency was compared to wtITR particles in a viable ex vivo human organ relevant to gene therapy applications, the cornea.^22^ Similar to the results of the mouse liver, SynITR vector transduction was increased over wtITR vectors up to 16-fold.

The collective work demonstrates that SynITR vectors were significantly enhanced for transduction compared to those harboring wtITRs, in a tissue-specific manner in a mouse model and in a human organ. Of note, in vitro transduction efficiencies of five different cell lines of variable tissue origin failed to reveal differences >0.7-fold between the wtITR and SynITR. This result highlights the importance of using animal models and viable human tissues to accurately assess the effects of SynITRs on the transduction efficiency. Additionally, wtITR vector transduction (but not SynITR) induced persistent γH2AX in the mouse liver and spleen, which is a marker of cellular stress, aging, and chronic inflammation.^23^ Taken together, SynITRs alter the vector-associated DNA damage response, and altered tissue-specific transduction, which could be exploited to develop low dose safer AAV gene therapies.

## Methods

### AAV vector production

All production plasmid herein relied on the standard 2 directional ITR sequences flanking the transgenic sequence context and relied on the EF1α promoter. The ITR sequence of serotype 2 (wtITR) or the synthetic ITR (SynITR)^10^ were used in these studies. Plasmid construction relied on molecular cloning and DNA synthesis and were maintained in SURE cells (Agilent, Santa Clara, CA). AAV vectors packaging EF1α-luciferase or EF1α expressing green fluorescent protein (GFP) with serotype 2 wtITR or SynITR^10^ were produced in HEK293 cells by the conventional triple transfection (pXX680 plasmid, pRepCap plasmid, and pITR plasmid) method.^24^ The AAV8-luciferase vectors used (in vitro assay, in vivo mice experiment, and ex vivo human cornea experiment) were produced by the UNC vector core and underwent iodixanol gradient purification followed by column chromatography purification and dialyzed in 350mM NaCl+5% D-Sorbitol in phosphate buffered saline (PBS; Table S1). The AAV2-GFP vectors used in the cell culture assays were produced by UNC vector core or in the Hirsch lab (TH and MH, Table S1) purified by cesium chloride ultra-centrifugation and subsequent dialysis in PBS.^24^

### AAV vector titer determination

Droplet digital PCR (ddPCR) was utilized for vector titering, crude lysate production analysis, and for the cDNA expression analysis. Vector titering was performed with TaqMan gene expression assays detecting GFP or EF1α sequences. Crude lysate production titering was performed with assays detecting sequences in the EF1α promoter (which was used in the production of all vectors) or transgene sequences (GFP, scIM^25^).

Before titering, vectors and crude lysates underwent DNase I (#M0303, NEB, Ipswich, MA) digestion in a reaction volume of 50µl containing 5µl each of vector, DNase I, DNase I reaction buffer (#B0303, NEB), pluronic F-68 (#24040032, Gibco) at 37°C for 30min, serial dilutions, and then capsid lysis at 95°C for 10min. ddPCR reaction was performed according to the manufacturer’s protocol using 2X ddPCR supermix for probes (No dUTP; #1863024, BioRad, Hercules, CA), a QX200 droplet generator (#1864002, BioRad), and analyzed by a QX200 droplet reader (#1864003, BioRad).

### Sequencing of the AAV capsid packaged genomes

The packaged genomes in AAV8-Syn/wt ITR-luciferase and AAV2-Syn/wt ITR-GFP were sequenced using Nanopore sequencing by Plasmidsaurus sequencing services (Eugene, OR).^26,27^ Raw sequencing reads were aligned to the reference viral genome using BWA-MEM on the UseGalaxy web interface.^28^ Alignments were visualized using Integrative Genome Viewer (version 2.19.4).^29^ Positions with high frequency mismatches (>20%) were highlighted in the coverage summaries (Fig. 1A).

**Figure 1.**
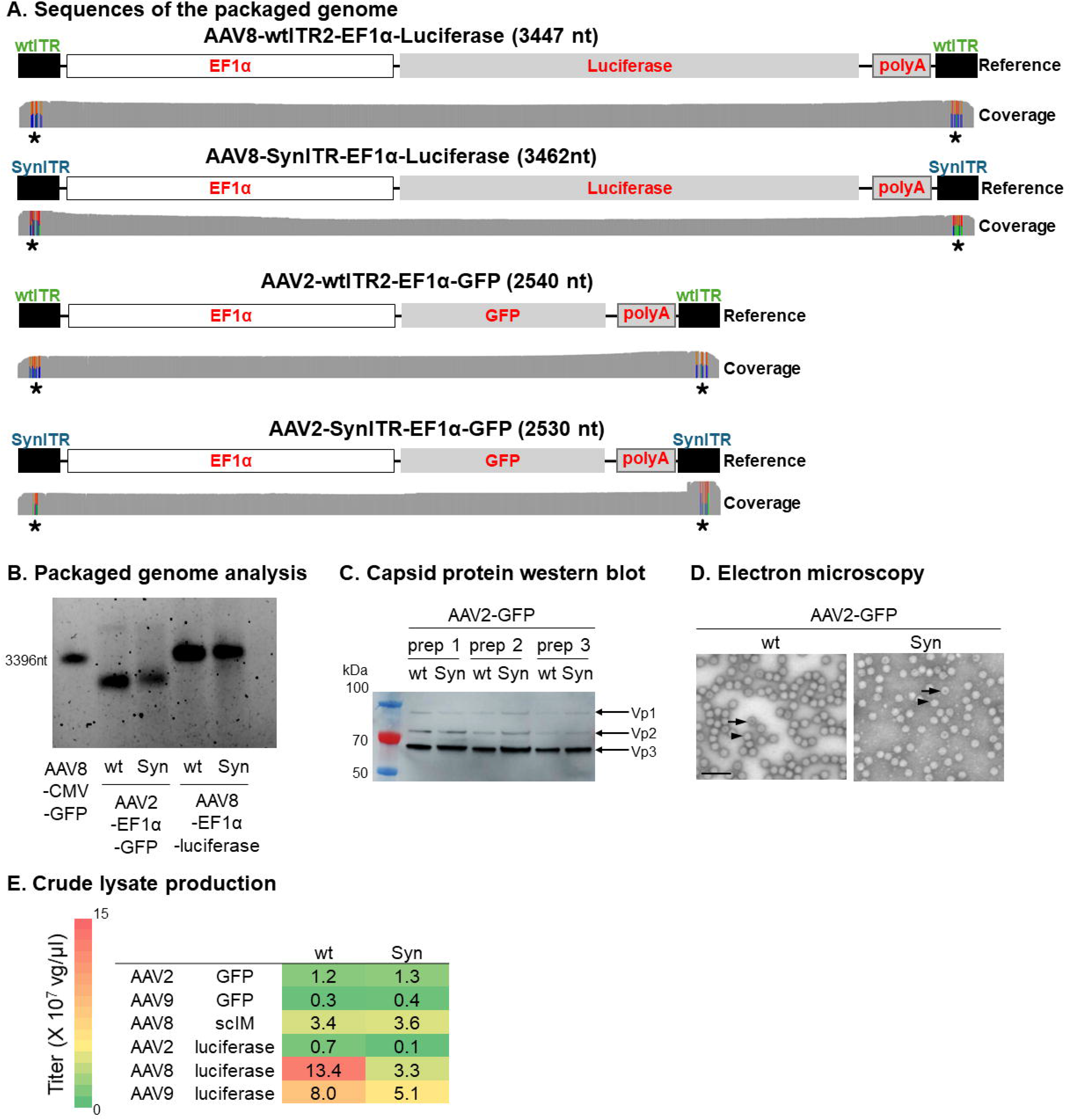
SynITRs produced equivalent vector preparations as wtITRs. **(A)** Sequence analysis of packaged vector genomes in adeno-associated virus serotype 2 (AAV2)-EF1α-wild type (wt) ITR-green fluorescent protein (GFP), AAV2-EF1α-Synthetic (Syn) ITR-GFP, AAV8-EF1α-wtITR-Luciferase, and AAV8-EF1α-SynITR-Luciferase aligned to an annotated reference genome. Coverage summaries represent the number of reads at each position, with >20% mismatches highlighted. Asterisks indicate regions of mismatches attributed to flip and flop orientations of the ITR. **(B)** Packaged genome analysis. Capsid packaged genomes (8.0e^9^vg) were analyzed by alkaline gel electrophoresis. A packaged wtITR vector genome (AAV8-CMV-GFP, 3396nt) was used as a size reference. **(C)** AAV capsid protein Western blot from preparations with wt or Syn ITRs. The same titer of the AAV2-GFP wtITR or SynITR vectors (8.0e8 vg/well) was used for analysis with an anti-AAV capsid 2 antibody. Three independent vector preparations (prep) were tested. **(D)** Electron microscopy images of AAV2-GFP wtITR or SynITR vector preparations. Black arrow heads and black arrows indicate full capsids and empty/partial capsids respectively. Scale bar: 100 nm. **(E)** Crude lysate vector production titers were assessed using ddPCR with an EF1α primer/probe set. The average titers of vectors with each capsid serotypes/transgenes with wtITR or SynITR are shown as a heatmap. N= 6each for AAV2-GFP, AAV9-GFP, AAV8-scIM, AAV2-luciferase and AAV8-luciferase, N=4 for AAV9-luciferase (Fig. S1C). scIM, single-chain immunomodulator.

### Alkaline gel electrophoresis

To determine the capsid packaged genome integrity, vectors were analyzed using alkaline gel electrophoresis as published.^10^ Briefly, 8.0e^9^ vg were digested with DNase I at 37°C for 1hr, followed by the addition of 0.5M EDTA to stop the reaction and then 10% sodium dodecyl sulfate to break open the capsid. The packaged genomes were denatured with 6X loading dye (0.4N NaOH, 5mM EDTA, 18% Ficoll and xylene cyanol),^24^ and run on an alkaline agarose gel (1% agarose gel containing 0.05N NaOH and 1mM EDTA) using alkaline running buffer (0.05N NaOH and 1mM EDTA) at 28V for 6hrs, and stained with SYBR Gold nucleic acid stain (#S11494, Thermo Scientific, Waltham, MA), followed by imaging with the Amersham ImageQuant 800 (Cytiva Life Sciences, Marlborough, MA).

### Analysis of viral protein incorporation in the AAV capsids

For western blotting of the purified vectors, samples were prepared with loading buffer (5% 2-mercaptoethanol in LDS sample Buffer, #NP0007, Invitrogen, Waltham, MA) and loaded onto a 4-12% BisTris gel (#NP0321BOX, Invitrogen) at an amount of 8.0e^9^ vg/well. Electrophoreses occurred in MOPS SDS running buffer (#NP0001, Invitrogen) and proteins were transferred to a nitrocellulose membrane using the iBlot2 system (#IB23002, Invitrogen). The membrane was blocked in 5% non-fat dry milk in TBST and subsequently incubated with an anti-AAV-Vp1/Vp2/Vp3 antibody (1:250, clone B1, ARP, #03-61058) followed by secondary antibody incubation and chemiluminescence imaging with the Amersham ImageQuant 800 (Cytiva Life Sciences). Three individually produced vectors (produced by TH, MH and UNC vector core) were tested (Table S1).

### Cell culture

HEK293 cells were cultured in Dulbecco’s modified Eagle’s medium (DMEM, #11995-065) supplemented with 4.5g/L of glucose, 10% bovine calf serum (BCS), and an antibiotic-antimycotic solution (#15240-062, Gibco). HepG2 human hepatocellular carcinoma derived cells were cultured in EMEM (#30-2003, ATCC, Manassas, VA) supplemented with 10% fetal bovine serum (FBS) and antibiotic-antimycotic. ARPE19 human retinal pigment epithelia cells were cultured in DMEM supplemented with 10% FBS with antibiotic-antimycotic. Highly proliferative human corneal endothelial cell line (HCEnC-21T)^30^ were cultured in medium containing 50% of Hams F12 Nutrient Mix (#11765-054, Gibco, Waltham, MA), 50% of Medium199 (#1150059, Gibco), supplemented with 5% FBS, 20 mg/L of ascorbic acid (#A4544, sigma, St. Louis, MO), 20 mg/L of insulin (#12585014, Thermo), 10µg/L of bFGF (#F0291, Sigma)^31^. 661W mouse photoreceptor derived cells were cultured in DMEM supplemented with 40 µg/L of hydrocortisone 21-hemisuccinate (#H-2270, Sigma), 40 µg/L of progesterone (#P-8783, Sigma), 32 mg/L of putrescine (#P-7505, Sigma), 40µl/L of 2-mercaptoethanol (#M-6250, Sigma), 10% FBS, and antibiotic-antimycotic. All cells were cultured at 37°C and in 5% CO_2_.

HepG2 cells were purchased from the UNC tissue culture facility. 661W cells were purchased from Dr. Muyyad R. Al Ubaidi.^32^ ARPE19 cells were provided by Dr. Zongchao Han^33^ and HCEnC-21T cells were supplied by Dr. Ula Jurkunas.^30^.

### Flow Cytometry

For transduction analysis using flow cytometry, cells were seeded in 24 well plates (HEK293 and HepG2 at 200,000 cells/well, ARPE-19 at 100,000 cells/well, HCEnC-21T at 60,000 cells/well, 661W at 40,000 cells/well) 24hrs prior to the addition of vectors. AAV2-GFP with wt or SynITR were added at 10,000 vg/cell for HEK293, HepG2, ARPE-19, HCEnC-21T cells and 50,000 vg/cell for 661W cells. After vector addition, cells were cultured for 48hrs (HEK293) or 72hrs (HepG2, ARPE-19, HCEnC-21T, and 661W cells) and then harvested for flow cytometry analysis. At harvest, cells were washed with PBS, detached with trypsin, washed with their respective culture medium, and re-suspended in PBS, then directly analyzed by flow cytometry. Propidium iodide (PI, #R37169, Invitrogen) was added to distinguish dead cells. Cells were analyzed with an Attune NxT instrument (Thermo). Forward scatter (FSC) and side-scatter (SSC) were used to segregate the target cell population, and after PI positive cells were gated out, the mean fluorescence intensity and the number of GFP positive cells were analyzed. The percentage of PI negative cells in transduced cells (GFP positive cells) was separately analyzed.

### Crude lysate AAV vector production assay

Production titers were assessed in crude lysates to allow rigor and to eliminate potential vector loss during purification methodology. HEK293 cells were seeded at 1,000,000 cells/well in 6 well plates 24hrs prior to transfection and were transfected with pXX680 (1.2 µg/well), pRep2Cap2 (0.8 µg/well), and pITR (0.8 µg/well) using polyethyleneimine (8.4 µl/well). Fifty-one hours after transfection, cell pellets were harvested, washed with PBS, re-suspended in 250µl PBS, and 3 freeze/thaw cycles were performed on the lysate alternating between a 37°C water bath and a dry ice bath. Lysates were clarified by centrifugation at 13,300rpm for 15mins, digested with DNase I, underwent capsid lysis, and the titers of crude lysate vectors were assessed by ddPCR using a Custom TaqMan primer probe assay targeting the EF1α sequence as described above.

### Experimental animals

This study was conducted in accordance with the policies on animal welfare of the National Institutes of Health and University of North Carolina (UNC). All protocols were approved by the Institutional Animal Care and Use Committee (IACUC) of UNC at Chapel Hill. Wild type C57BL/6J mice were kept at the Division of Comparative Medicine facility at UNC and fed ad libitum. Mice were monitored 3 times a week to check the body weight and body condition score.^34^ The criteria for euthanasia before the end of the study were set to the UNC standard of humane endpoints (including greater than 20% total weight loss and body condition score of 2).

### Intravenous AAV vector injection and luciferase imaging in mice

Six week old C57BL/6J male mice were randomized by weight into study groups (AAV8-EF1α-luciferase vectors with wt or Syn ITRs) and were administered 8.0e^10^vg in 100µl of the assigned vector by intravenous tail vein injections. Non-injected mice served as non-treated controls (NT). All tail vein injections were performed by a single experienced researcher at the UNC Preclinical Research Unit.

The mice were subjected to live luciferase imaging on days 3, 7, 14 and 21 post-injection while under anesthesia by inhalation of isoflurane (1 – 4% isoflurane inhalation). Luciferin (125µl at 25 mg/ml) was intraperitoneally injected 5mins before imaging. Imaging was performed using the IVIS-Lumina (Revvity, Inc., Waltham, MA) and analyzed using Living Image v4.7.4. software (Revvity).

On day 23 post-injection, euthanasia was induced by CO_2_ compressed carbon dioxide gas followed by cervical dislocation and the liver, heart, tibialis anterior (TA) muscles, kidneys, spleen and pancreas were collected from each animal and frozen immediately.

All animal handlers and researchers who performed the procedures and analysis were blinded to the group assignment throughout the in-life and post-mortem analyses.

### Ex vivo human cornea

Human donor corneas were obtained from the Miracles In Sight eye bank. The corneas were evaluated at Miracles In Sight and deemed eligible for research use. Corneas of both eyes from the same donor were obtained. The criteria to be included in this study were the following: post-harvest to injection time could not exceed 6 days, no current history of sepsis, not currently under treatment with chemotherapy, no moderate to severe corneal stromal infiltration, and eligibility of both corneas from the same individual. The cornea was excised at Miracles In Sight and stored in Optisol-GS (Bausch+Lomb, #50006-OPT, Laval, Quebec, Canada) and shipped on ice. After delivery, the two corneas from each individual were randomly assigned to the experimental groups and 1.0e^10^vg (50µl)^35^ of AAV8-EF1α-luciferase vectors with wt or Syn ITRs were injected to the corneal stroma using a 31 G needle by a single researcher. The corneas were cultured in Optisol-GS at 37°C for 7 days unless otherwise noted.

The researcher who performed all procedures and analysis was blinded to the group assignment throughout the procedures and analyses.

### Luciferase assay (HEK293 cell, mice tissues, human corneas)

HEK293 cells were seeded at 200,000 cells/well into 24 well plates 24hrs prior to the addition of AAV8-luciferase vectors with wt or Syn ITR at 2,500 or 10,000 vg/cell. After 72hrs, cells were washed with PBS, harvested with 100µl of passive lysis buffer (#E194A, Promega, Madison, WI) followed by incubation for 5mins at 23°C with shaking at 140rpm, thoroughly vortexed, and centrifuged at 13,300rpm for 5mins at 4°C.

The frozen mouse tissues were homogenized, and 40mg of each tissue was harvested in 400µl of passive lysis buffer containing Halt protease inhibitor (#78430, Thermo Scientific), followed by incubation for 30mins at 23°C with shaking at 1200rpm and centrifugation at 13,300rpm for 5mins at 4°C.

Human corneas were washed with cold PBS before harvest and the epithelia was removed by blade followed by another wash with PBS. After excision of the scleral ring, the corneas were homogenized and 900µl of passive lysis buffer with Halt protease inhibitor was added. The mixture was thoroughly vortexed and subsequently incubated at 23°C for 45mins with shaking at 1200rpm with occasional vortexing during incubation, followed by another thorough vortex, and the lysate was harvested after centrifugation at 13,300rpm for 40mins at 4°C.

The luciferase activities of the protein samples were analyzed using 20µl of sample for each cell lysate or human cornea samples, or 100µl of mouse tissue samples with 100µl of reagent (#E1483, Promega) in a 96 well plate using the Glomax plate reader (Promega). The luciferase activity results were normalized to the total protein concentration (Bio-Rad Protein Assay, #5000006, BioRad). For protein concentration quantification, 200µl of protein assay dye was added to 10µl of sample on a 96 well plate and the protein concentration was calculated from the absorbance measured at 562 nm using the BioTek Cytation 5 plate reader (Agilent, Santa Clara, CA).

### Western blotting mice tissues

Mouse tissue (liver, heart, muscle, kidney, pancreas, and spleen) samples harvested in passive lysis buffer (same samples used in the luciferase assay above) were analyzed by western blotting as described above in the virus western blotting section. For mouse tissues, 50µg of total protein was loaded. Anti-phospho-histone H2AX (Ser 139) (#05-636, clone JBW301, Milipore Sigma, Bedford, MA), anti-phospho-p53 (Ser 15, #12571, clone D4S1H, Cell Signaling, Danvers, MA), and anti-β-actin (#sc-47778, clone C4, Santa Cruz Biotechnology, Dallas, Texas) were used for the reaction. For phospho-p53 analysis, etoposide treated human fibroblasts which expresses phospho-p53^1^ was used as a positive control. The γ-H2AX band intensities of each sample were measured using ImageJ ^36^ and were normalized to β-actin band intensities.

### DNA and cDNA preparations for mice tissues

Both DNA and mRNA were extracted simultaneously from homogenized mouse tissues (20mg for liver and spleen, and 50mg for heart, TA muscle, kidney and pancreas) with the Zymo Quick-DNA/RNA Miniprep Plus Kit (#D7003, Zymo Research, Irvine, CA) according to the manufacturer’s protocol and eluted in 100μL of nuclease free water. Concentrations and quality of the DNA and RNA were determined by NanoDrop One (Thermo Scientific) and agarose gel electrophoresis, respectively. A DNase Digest was performed on all mRNA samples with the Invitrogen TURBO DNA-free Kit (37°C for 30 min, #AM1907, Thermo Scientific) followed by cDNA synthesis using the High Capacity cDNA Reverse Transcription Kit (#4368814, Thermo Scientific) according to the manufacturer’s protocol.

### Vector genome biodistribution analysis for mice tissue

Quantitative reverse transcription polymerase chain reaction (qPCR) was utilized for vector genome analysis in mouse tissues. DNA extracted from mouse tissues were analyzed by qPCR using TaqMan Universal PCR Master Mix (#4304437, Thermo Scientific) and a custom TaqMan Gene expression assay for the EF1α sequence (as described below) with the StepOne Plus thermocycler (Applied Biosystems) at an annealing temperature of 55.8°C. A CT value of ≤ 35 cycles was set as the detection limit.

### Transcript expression analysis

Transcript expression analysis was performed on cDNA from mouse liver samples using TaqMan gene expression assays detecting cyclin-dependent kinase inhibitor 1A (p21), BCL2 associated X apoptosis regulator (Bax), and hypoxanthine phosphoribosyltransferase 1 (Hrpt1) as a housekeeping control.

The following TaqMan gene expression assays were used for detection; Custom assay (#4351372, Thermo Scientific), EF1α_forward; GGAAGTGGGTGGGAGAGTTC, Reverse; GAAGGTGCCACCAGATTCG, probe; CGCCCAGGCCAGGCCTCAAC, scIM_forward; ATCCTGCCGAGATCATCCTG, Reverse; GTGTACCTCTGCTCCTCTCC, probe; ACCTGCCGGCGATGGCACCT, Inventoried assays (#4331182, Thermo Scientific) GFP; Mr04329676mr, p21; Mm01303209_m1, Bax; Mm00432051_m1, Hrpt; Mm03024075_m1. Annealing temperature used was 55.8°C for EF1α, 55.9°C for scIM, 60°C for all the inventoried assays.

### Statistics

Head-to-head comparisons between wt and Syn ITRs were performed with an unpaired t-test (crude lysate titers, transduction in vitro with flow cytometry). Multiple comparisons were performed with a one-way analysis of variance (ANOVA) with Tukey’s multiple comparisons test (transduction efficiency in vitro with luciferase, in vivo postmortem luciferase assay) or with a two-way ANOVA with Tukey’s multiple comparisons test (transduction efficiency in vivo in life over time analysis). The level of statistical significance was set at p < 0.05.

## Results

### SynITR AAV vector preparations were equivalent to those generated with wtITRs

To determine whether AAV vectors packaged with SynITRs were comparable to wtITR AAV vectors, characterization of viral particles was performed using multiple capsid serotypes (AAV2, AAV8, and AAV9) and three different transgenes: luciferase, GFP, and the single chain immunomodulator, recently reported to prevent allogenic corneal transplant rejection (scIM).^25^ First, to confirm the SynITRs were packaged by natural AAV capsids, AAV8-Syn/wt ITR-luciferase and AAV2-Syn/wt ITR-GFP packaged genomes were analyzed via Nanopore sequencing. The results demonstrated that the SynITR nucleotide sequence is packaged in the flip/flop orientations similar to wtITR sequence (Fig. 1A). Additionally, the majority of capsid particles contained the full-length cassette regardless of the serotype (Fig. 1A). Next, the packaged vector genome integrity was investigated by alkaline gel electrophoresis. The data demonstrated that similar DNA species corresponding to the intended packaged genome sizes were packaged in the AAV2 or AAV8 capsids regardless of the wt or Syn ITR sequence (Figs. 1B, S1A). Furthermore, western blotting of capsid proteins loaded at equal vector genomes (vg) of three different AAV2-GFP preparations showed comparable amounts of capsid proteins (Vp1, Vp2, Vp3) at their expected ratios in vectors generated with Syn or wt ITRs (Fig.1C). This result also suggests a similar full/empty capsid packaging ratio which was further supported by >94% full capsid packaging as determined via electron microscopy (Fig. 1D, Suppl. Fig. S1B, Table S1). The final production titers were determined by ddPCR with multiple primer sets. SynITRs generally produced equivalent or elevated titers compared to wtITRs independent of the capsid serotype (AAV2, 8, or 9) when packaging the GFP or scIM transgene^25^ (Figs. 1E, S1C). In contrast, when the luciferase gene was packaged, wtITRs produced significantly higher titers than SynITRs in all capsids tested (AAV2, 8, or 9), even though the SynITR luciferase titers were comparable to vectors produced with the other transgenes. (Fig. 1E, Table S1). Taken together, these data demonstrate that SynITRs produce AAV vectors at similar titers and quality as wtITRs and variations in production titers are noted amongst transgenes and capsids for each ITR sequence.

### AAV-SynITR vectors are decreased for transduction compared to wtITRs in cultured cells

To investigate whether the SynITRs altered transduction efficiency in cultured cells, wt and Syn ITR AAV2-GFP vectors were evaluated in five cell lines of various species and tissue origins including the liver (HepG2), retina (661W and ARPE19), cornea (HCEnT-21T), and kidney (HEK293). In all of the five tested cell lines, the transduction efficiency of vectors harboring the wtITRs was modestly higher (<10%) than in cells transduced with the SynITR vectors (Figs. 2A - F, S2A). To investigate whether the decrease in transduction efficiency mapped to the SynITR sequence, AAV8-luciferase vectors were evaluated in HEK293 cells (Fig. 2G). At the same vg/cell ratio with the AAV2-GFP context (10,000 vg/cell), the SynITRs showed a 21.8% decrease in transduction compared to wtITRs while at a dose of 2,500 vg/cella similar decrease of 23.4% was observed (Fig. 2G). Cell viability was indirectly assessed using PI staining and, in all cases, remained unchanged following transduction with wtITR or SynITR AAV2 vectors (Fig. S2B).

**Figure 2.**
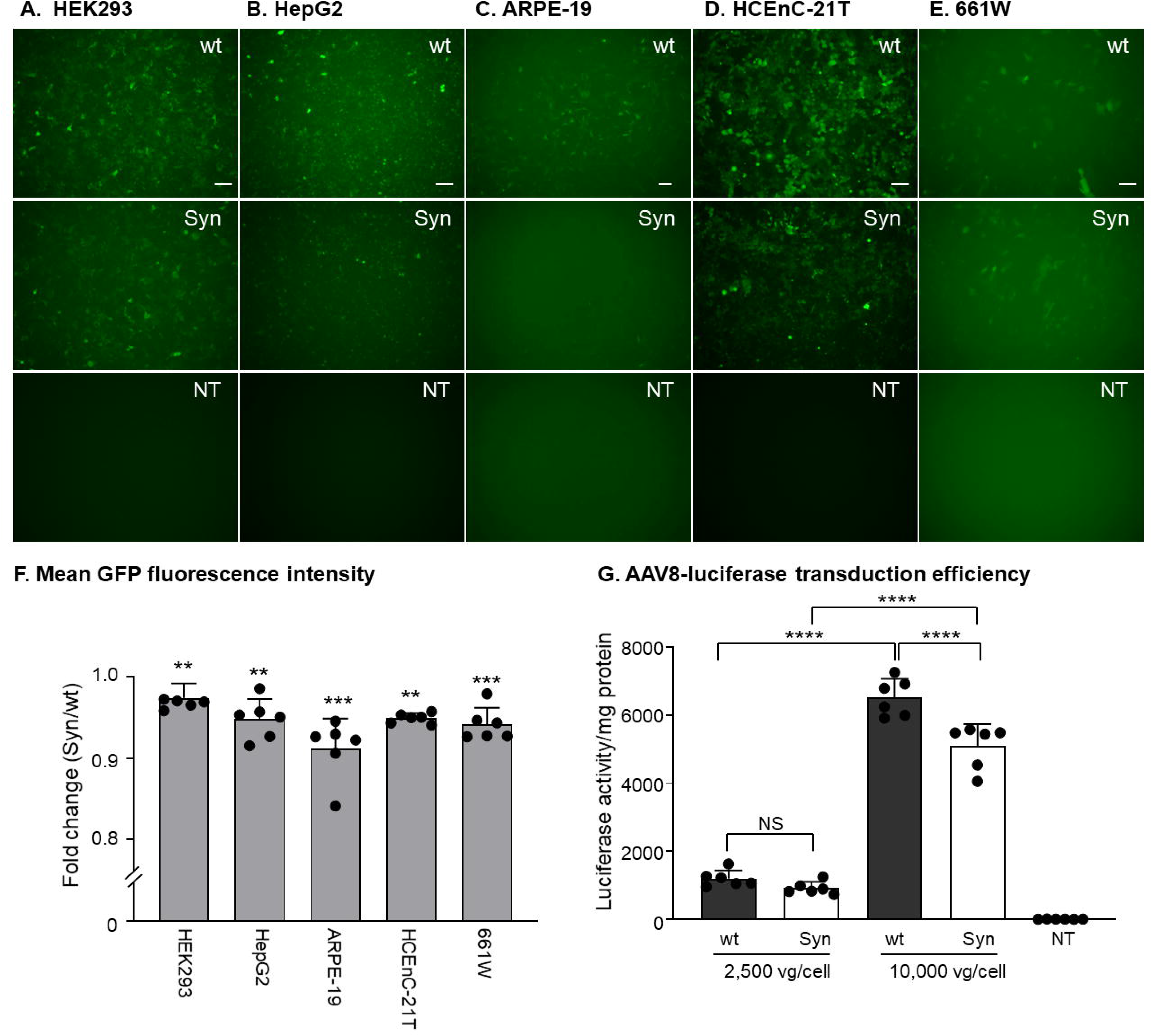
AAV SynITR exhibited diminished transduction efficiency in cultured cells. **(A-F)** human embryonic kidney (HEK) 293 cells (**A**), HepG2 human hepatocellular carcinoma derived cells (**B**), adult retinal pigment epithelium (ARPE)-19 cells (**C**), highly proliferative human corneal endothelial cell line (HCEnT-21T) (**D**), or 661W mouse photoreceptor derived cells (**E**) were transduced with AAV2-GFP with wt or Syn ITRs at a vector genome (vg) to cell ratio of 10,000 for HEK 293, HepG2, ARPE-19, and HCEnT-21T and 50,000 vg/cell for 661W cells. After vector addition, cells were cultured for 48hrs (HEK 293) or 72hrs (HepG2, ARPE-19, HCEnT-21T, and 661W) before flow cytometry analysis. Scale bars: 100 µm in (**A**, **B**, **D**, and **E**), 200 µm in (**C**). (**F**) Transduction efficiency measured by flow cytometry, mean intensity is shown as fold change of Syn vs wt ITRs. The fold change was calculated by dividing value of each sample by the average of wtITR samples. N=6 each, **p<0.01, ***p<0.001, unpaired t-test. Data are presented as mean ± standard deviation. **(G)** AAV8-luciferase transduction efficiency. HEK 293 cells were transduced with AAV8-luciferase vectors with wt or Syn ITRs at 2,500 or 10,000 vg/cell. Luciferase activity was measured in cell lysates harvested 72hrs after vector addition, normalized to the total protein concentration, and shown as luciferase activity/mg total protein. N=6 each, ****p<0.0001, NS, not statistically significant, Tukey’s multiple comparison test. Data are presented as mean ± standard deviation. All of the infected cells showed significant luciferase activity compared to NT; p<0.01 vs. SynITR at 2,500 vg/cell, p<0.001 vs. wtITR at 2,500 vg/cell, and p<0.0001 vs. wt and Syn ITRs at 10,000 vg/cell. NT=no transduction.

### SynITRs enhanced transduction in vivo in a tissue-specific manner without altering vector biodistribution when compared to wtITRs

Next, to determine if the SynITR altered transduction in vivo, AAV8-EF1α-luciferase with Syn or wt ITRs were administered by tail vein injection to male C57/BL6J mice. The same vector preparations characterized in Fig. 1 and evaluated in the cell culture study (Fig. 2G) were used at a dose of 8.0e^10^vg in a final volume of 100µl and non-injected mice served as the negative control. No animal in any group exhibited weight loss, nor were there any signs of distress observed throughout the study (Fig. S3A). The body condition score^34^ of all animals remained at 3 throughout the study period. In vivo detection of luciferase activity from both Syn and wt ITR AAV8 vectors demonstrated similar levels of signal that were significant compared to the non-injected control mice as early as day 3 post-injection (Fig. 3A, B). At all time points thereafter (days 7-21), in-life imaging demonstrated that the SynITR vector exhibited up to a 4.0-fold enhancement of luciferase activity compared to the wtITR vector in the abdomen (p<0.05, p<0.001, and p<0.01 at day 7, 14, and 21 respectively, Tukey’s multiple comparison test, Fig. 3B).

**Figure 3.**
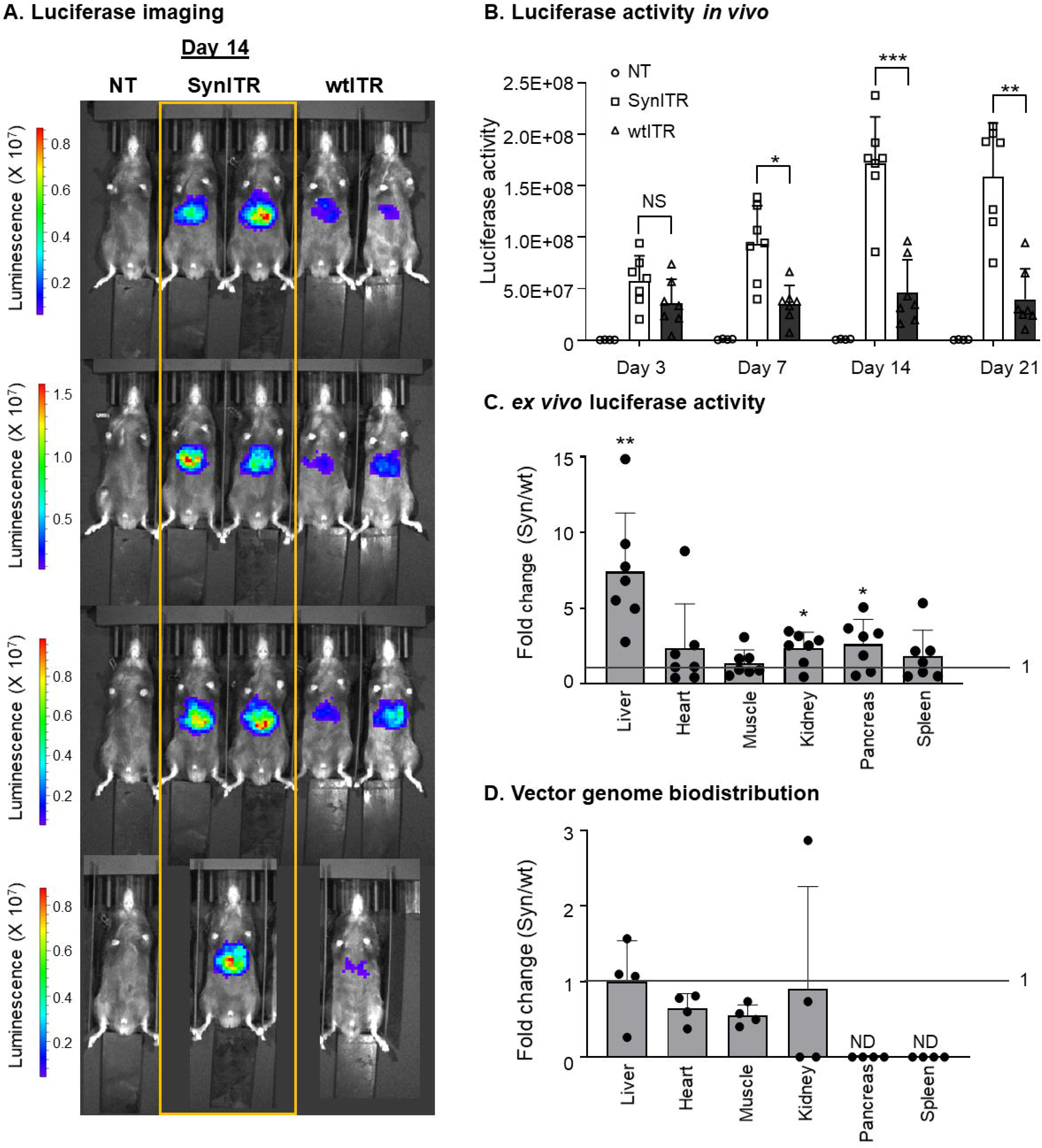
SynITR vectors exhibit enhanced tissue specific transduction. Wild type (wt) six week old male mice were administered 8.0e^10^vg of AAV8-EF1α-luciferase harboring wt or Syn ITRs by tail vein injection. Non-injected mice served as non-treated controls (NT). Mice underwent in vivo imaging on days 3, 7, 14, and 21 post-injections. **(A)** In vivo luciferase activity imaged 14 days after vector injections. Luminescence intensities are shown on a color scale of radiance (p/sec/cm^2^/sr). **(B)** In vivo luciferase activity over time shown as total flux (p/sec). N=7 for wt and Syn ITR vector injected mice, N=4 for NT. *p<0.05, **p<0.01, ***p<0.001, NS=not statistically significant, Tukey’s multiple comparison test. Vector injected mice (either wt or Syn) showed statistically significant luciferase activity at all examined time points compared to NT, at day 3, p<0.01 vs SynITR and p<0.05 vs wtITR, at day 7, p<0.01 vs SynITR and wtITR, at day 14 and day 21, p<0.001 vs SynITR and p<0.05 vs wtITR. Data were presented as mean ± standard deviation. **(C)** Ex vivo luciferase activity in homogenized whole organs. Liver, heart, muscle (tibialis anterior muscle), kidney, pancreas, and spleen were collected from mice 23 days post-injections. The luciferase activity was normalized to total protein concentration and is shown as a fold change of Syn vs wt ITR. N=7 for wt and Syn ITR vector injected mice, N=5 for NT. *p<0.05, **p<0.01, unpaired t-test. Data are presented as mean ± standard deviation. **(D)** Vector genome biodistribution shown as a fold change of Syn vs wt ITR. Vector genome copy number was analyzed by quantitative polymerase chain reaction (qPCR). ND = not detected in all samples. The fold change was calculated by dividing the value of each sample by the average of wtITR samples. N=7 for wt and Syn ITR vector injected mice, N=5 for NT. Data are presented as mean ± standard deviation.

On day 23 post-injection multiple tissues were dissected and frozen for additional analysis. Consistent with the in-life imaging analysis, a postmortem luciferase assay using homogenized liver samples demonstrated a 7-fold enhancement of luciferase activity in animals treated with the SynITR vector compared to the wtITR vector (p<0.01, t-test, Figs. 3C, S3B). Furthermore, post-mortem homogenized kidney and pancreas samples also showed enhancements of luciferase activity of 2.3 and 2.6-fold respectively, in SynITR vector treated subjects compared to the wtITR vector (Fig. 3C). In contrast, no significant difference in luciferase activity was found between wt or Syn ITR vectors in homogenized heart, skeletal muscle, or spleen lysates (Figs. 3C, S3B). To determine whether SynITR vectors exhibited enhanced luciferase activity simply due to an altered biodistribution or stability, vector genomes were quantified from various tissues. The data revealed similar vector genome copy numbers were present in both the wt and Syn ITR treated animals in all tissues tested, suggesting that the SynITR did not affect particle tissue targeting or intracellular vector genome stability throughout the duration of this study (Figs. 3D, S3C).

### wtITRs, but not SynITRs, induced persistent γH2AX in the liver and spleen

It has been extensively published that wtAAV vector transduction induces DNA damage signaling,^37–40^ and more recently that SynITR vector transduction alters this response, at least in hESCs.^10^ To investigate whether the SynITRs altered the DNA damage response in vivo, p53 signaling was investigated in the liver 23 days post-injection. At this timepoint, p53 activation by phosphorylation of Ser15 (an early marker of DNA damage signaling) was not observed in the liver samples of either the SynITR or wtITR treated groups (Fig. 4A). Consistently, expression of p53 transactivation targets p21 (associated with G1/S phase cell cycle arrest) and Bax (associated with apoptosis) were not significantly altered among the non-treated, wtITR, or SynITR experimental groups (Fig. 4B, C).^41,42^ Next, gamma-H2AX (γH2AX), another early marker of DNA damage and aging that is induced by the wtITR sequence was investigated by western blotting^18,23,43^. Surprisingly, at day 23 post-injection γH2AX was abundant in the spleen and liver, but not muscle, heart, pancreas, or kidney, of all six wtITR vector treated mice (Figs. 4D, E, S3D, S3E). In contrast, minimal to no γH2AX was observed in any of the SynITR vector transduced tissues (Figs. 4D, E, S3D, S3E).

**Figure 4.**
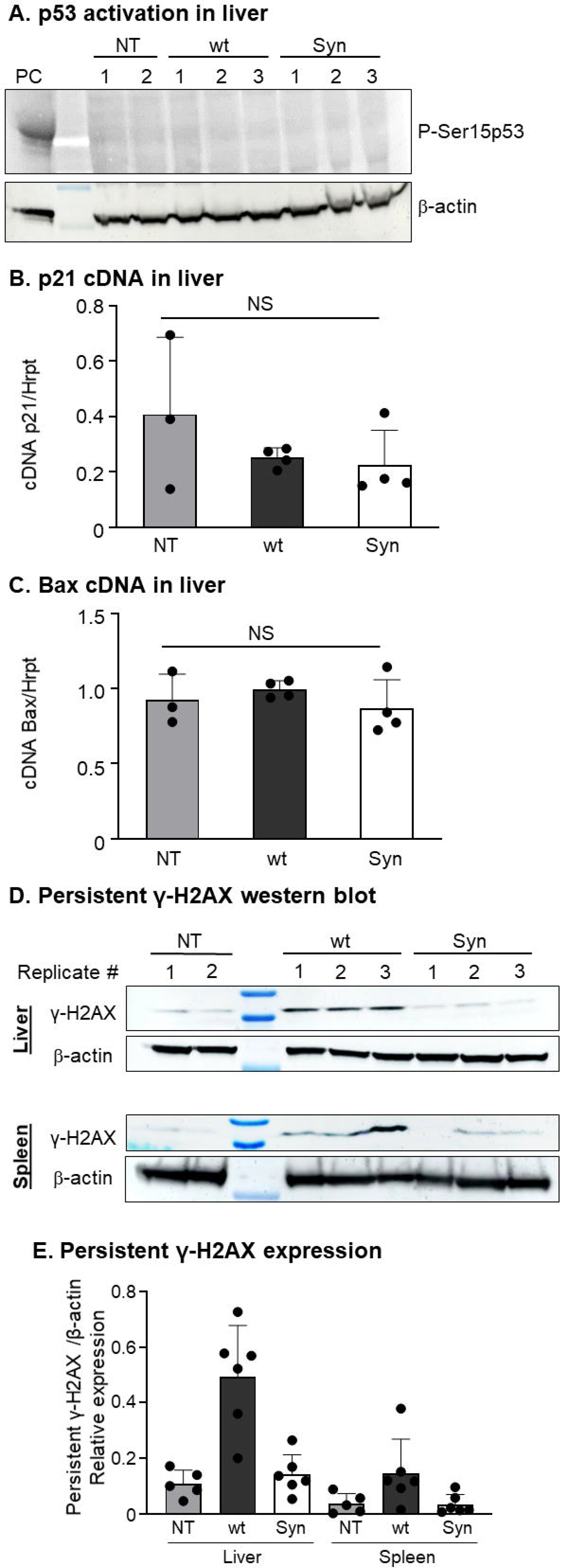
p53 and γ-H2AX activation following wt or Syn ITR vector transduction in mice tissues. Wild type six week old male mice were administered with 8.0e^10^vg of AAV8-EF1α-luciferase vectors harboring wt or Syn ITRs by tail vein injections. Non-injected mice served as non-treated controls (NT). Liver, spleen, kidney, pancreas, heart and muscle (tibialis anterior muscle) were collected from mice 23 days post-injection. **(A)** Western blot analysis of phospho-p53 (Ser15) in mouse liver. An etoposide treated human fibroblasts sample was used as a positive control (PC). β-actin was used as a loading control. N=2 mice for NT, N=3 mice each for wt and Syn ITR. **(B, C)** The cDNA of p53 activated transcripts cyclin-dependent kinase inhibitor 1A (p21, **B**) and BCL2 associated X, apoptosis regulator (Bax, **C**) in mouse liver were analyzed by ddPCR. The cDNA level was normalized to Hrpt1 cDNA. N=4 animal each. NS, not statistically significant, Tukey’s multiple comparison test. Data are presented as mean ± standard deviation. **(D, E)** Western blot analysis of phospho-histone H2AX (γ-H2AX; Ser139) of mice liver, and spleen. The same replicate number in each treatment group represents the sample from the same animal in (**A**), (**D**) and (**S3D**). Quantification of γ-H2AX band intensity was normalized to β-actin **(E)**. N=5 mice for NT, N=6 mice each for wt and Syn ITRs (Fig. **S3E**). Data are presented as mean ± standard deviation.

### SynITRs increased transduction in human corneas compared to wtITRs

To determine whether the SynITR transduction enhancement in particular mouse tissues translates to human tissue, a therapeutically relevant cornea model^25^ measuring ex vivo rAAV transduction was performed. Briefly, human donor corneas from 4 individual male patients were injected with AAV8-luciferase vectors containing the SynITR in one cornea and the wtITR in the contralateral cornea to control for inter-individual genetic variation. The intrastromal injections (50µl) were performed without perforations or any major complications and harvested 5-7 days post-injection (Fig. 5A). All corneas injected with either vector showed elevated levels of luciferase activity compared to the PBS injected vehicle control cornea (Fig. 5B, 5C). Similar to the results observed in the liver, kidney, and pancreas for the IV-injected mice, all human corneas injected with the SynITR vector exhibited enhanced luciferase activity up to 16-fold (2.6-fold to 16-fold with a median of 7.7-fold) compared to the contralateral wtITR vector injected corneas (Fig. 5B, 5C).

**Figure 5.**
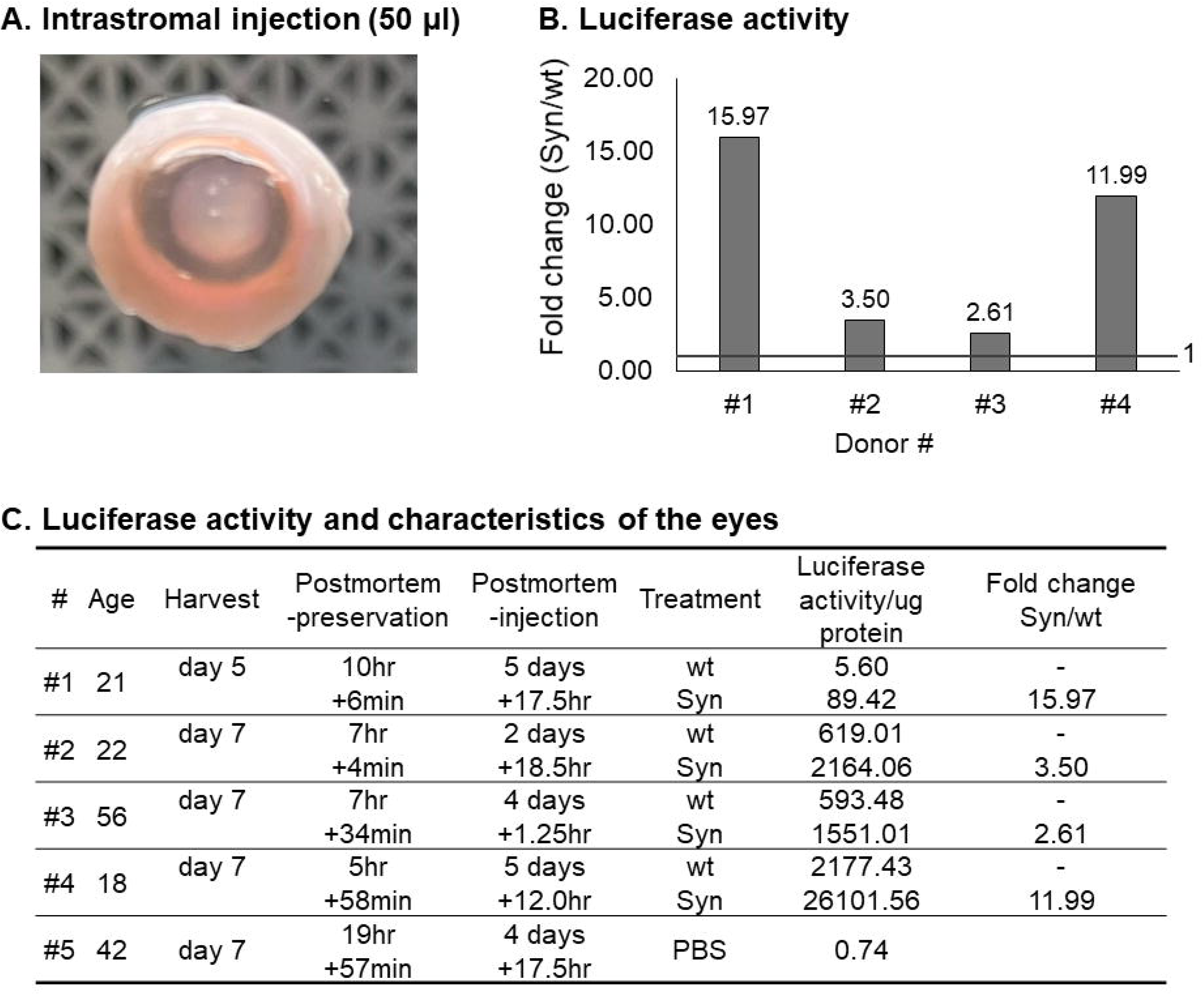
SynITRs enhanced vector transduction in human corneas ex vivo. AAV8-EF1α-luciferase vectors with wt or Syn ITRs were injected to the stroma of human donor corneas at a dose of 1.0e^10^vg in 50µl. **(A)** The two corneas from the same individual were randomly assigned to wt or Syn ITR vectors. After culturing for 5-7 days, luciferase activity of the protein harvested from the corneas was measured. **(B)** Luciferase activity of the corneas is shown as a fold change of Syn vs wt ITR. Fold change was calculated as a comparison within the corneas from the same individual. N=4 individuals. **(C)** Summary of luciferase activity and the demography of the donor corneas. PBS=phosphate buffered saline.

## Discussion

AAV vectors are commonly used in gene therapy for a variety of diseases in different targeted tissues; however, off-target expression, vector-associated toxicities, relatively poor transduction efficiencies, and high vector production costs limit widespread applications^1–3,44,45^. In a previous report, a SynITR ablated for p53 binding sites inherent to wtITR altered AAV vector-induced DNA damage signaling and attenuated apoptosis in hESCs.^10^ While the previous report used vectors originating from the DD plasmid context, which packages a bacterial replication origin and antibiotic resistance gene^10,46^, herein AAV vectors were produced from a standard 2 ITR (each with a single D) plasmid context containing the SynITR sequence,^10^ and were tested for vector production and transduction efficiency in vitro, in vivo, and ex vivo. AAV vectors harboring SynITRs demonstrated cell type specific decreased transduction in cultured cells and, in contrast, increased transduction in vivo in a tissue-specific manner. When systemically administered to wt C57/BL6J mice, SynITR significantly enhanced transduction in the liver, kidney, and pancreas at similar vector copy numbers. Interestingly, persistent γH2AX, a marker of cellular senescence and chronic inflammation,^43,47^ was abundant in the liver and spleen harvested from mice administered the wtITR, but not the SynITR, vector. Moreover, SynITR was tested in human corneas, and enhanced transduction 2.6-fold to 16.0-fold compared to wtITRs, perhaps highlighting the trans-species conserved functionality of SynITRs. These collective SynITR data emphasize the potential for expansion to clinical applications towards low-dose, safer, and tissue-specific gene therapy. Additionally, the SynITRs offer an alternative vector context capable of high titer production to elucidate wtITR biology including the tissue-specific DNA damage response elicited upon transduction.

One barrier to the widespread use of AAV vectors is the cost of high titer preparations.^44^ Production titers can vary depending on the capsid used for packaging^48^ or the transgenic sequence.^49^ Because the ITR serves as the AAV replication origin,^9^ changing its sequence could theoretically affect the replication efficiency and, consequently, the overall titer. However, when originating from the DD plasmid context, the SynITR was not significantly altered for replication when compared to the wtITR yet was significantly increased for production approximately 3-fold.^10,46^ From a 2 ITR standard AAV plasmid context, the SynITR produced similar titers compared to wtITR in a manner mostly independent of capsid serotypes, except that SynITR produced higher titers when GFP was packaged in the AAV9 capsid (Fig. 1E, S1C, Table S1). The production titers were partially independent of transgenic DNA, however, specifically the luciferase gene resulted in higher titers with the wtITRs in a capsid-independent manner (Fig. 1E, S1C). While the transgene itself can affect the production titer,^49^ the size of the packaged gene located between the ITRs can also affect the packaging.^50^ In the current study, the size of transgene cassettes between the ITRs were 2540nt (EF1α-GFP), 3447nt (EF1α-luciferase), and 3576nt (EF1α-scIM) all under the <5kb size constraint of AAV capsids. Furthermore, packaged genomes were confirmed to have a primary single DNA species at the expected sizes which was supported by genome sequencing (Figs. 1, S1A). Interestingly, the high titer production using the luciferase cassette appears to be capsid-dependent as both wt and SynITR titers increased over 10-fold using serotypes 8 and 9 compared to serotype 2, an effect that was not observed for the GFP cassette regardless of the ITR sequence (Fig. 1E). Notably, the production titers were consistent using two different purification methodologies and in unpurified lysates (Figs. 1, S1, Table S1). Additionally, the SynITRs not only produced similar titers to wtITR, but also produced vectors with comparable full/empty capsid ratios as determined by distinct assays (Fig. 1C, D). Most importantly, the sequencing results clearly demonstrate the capsid packaging of the SynITRs in the flip/flop orientations similar to the packaged ssDNA species of wtAAV and replicated wtAAV genomes (Fig. 1A)^51,52^

Other barriers for AAV gene therapy applications include off-target expression, liver and neurological toxicities, and relatively poor transduction efficiencies.^2,44,53^ In order to achieve low dose gene therapy with reduced undesirable side effects, enhanced transduction in target tissue is actively pursued largely at the levels of the capsid and promoter. Herein, the rationally designed SynITR was tested to serve this purpose in cell culture, in vivo, and in human tissue ex vivo. Interestingly, even though the SynITR transduction was modestly diminished <10% in all cell lines tested, with a minimal dose effect (Fig. 2), the SynITR enhanced transduction in vivo up to 7-fold compared to wtITR (Fig. 3). The transduction enhancement was tissue-specific; enhanced significantly in liver, pancreas, and kidney but not in the heart, muscle, and spleen. Previous reports had shown that while IV administered vectors accumulate the most in liver tissue, the biodistribution of the vectors in spleen, heart, and muscle was similar to that in the pancreas and kidney, where SynITR showed enhanced transduction.^54,55^ Notably, the vector genome copy number was similar between SynITR and wtITR in all tissues tested, even in the tissues where SynITR showed enhanced transduction (Fig. 3). Thus, the tissue-specific transduction by SynITR is unlikely due to the altered vector genome biodistribution. In some of the tissues (pancreas and spleen) vector genomes were at or near the limit of detection, which may be a consequence of inefficient tissue lysis and the highly vascular nature of spleen, which can cause carryover of PCR inhibitors such as hemoglobin or immunoglobulin G.^56,57^ Another possible factor for tissue-specific enhanced transduction by the SynITR is the cell types and cycling conditions including varied transcription factor expression. For example, the transduction efficiency by wt or SynITRs were variable in cell lines of different origins (Fig. 2). Furthermore, in vivo, cells in some organs are terminally differentiated or quiescent (e.g. cardiomyocytes in adults and muscle fibers) while others have the capacity to divide (e.g. hepatocytes, proximal tubule cells in the kidney, β-cells in the pancreas, and germinal center cells in the spleen).^58–60^ At the test article level, the SynITR produced vectors of comparable quality to wtITR vectors (including full/empty ratio and full length cassette packaging) which decreases concerns of such aspects altering the transduction efficiency (Figs. 1, S1). Thus, other possible factors for transduction enhancement by the SynITR may be exerted at any step of the rAAV infection including differential uncoating, second-strand synthesis, chromosome integration, episome formation/conformation kinetics and/or epigenetic modulation.^61^ A bioinformatics analysis of the putative transcriptional factor binding sites inherent to the wtITR, including p53 sites, revealed that SynITR is varied for predictions in >20 factors.^10^ It is possible that different transcription factors which bind to SynITR, but not to wtITR, or vice versa, affect the enhanced transduction such as repression of the wtITR vector genome by the epigenetic regulator γH2AX. This may be related to the unrepaired DNA double strand breaks at human telomeres which often exhibit persistent γH2AX coupled with γH2AX’s general role in epigenetic repression. ^23^ However, this hypothesis remains to be formally validated. Since wtITR rAAV transduction efficiency is influenced by sex, ^62,63^ a limitation of this current study is that the in vivo and ex vivo transduction experiments were conducted using only male mice and corneas. Investigations to determine if SynITR transduction enhancement is biased upon sex are in progress.

AAV has previously been shown to cause a host cell DNA damage response upon transduction,^18,39,47^ and in particular cell types, such as hESCs, it induces p53-dependent apoptosis.^1,38,64–66^ The SynITR used herein was rationally designed to ablate the p53 binding sites to attenuate the induced apoptotic signaling pathway in hESCs.^10^ When livers from treated mice were harvested and tested, p53 signaling was not detected either in wtITR or SynITR treated mice (Fig. 4). This may be due to the timing of the harvest (day 23 post-injection), as p53 activation is typically a rapid response occurring within several minutes to hours post-treatment.^67–70^ Interestingly, from the same mouse tissues, the abundance of persistent γH2AX, a DNA damage response marker which plays a multitude of roles in DNA repair, chromatin remodeling, cellular senescence, and gene silencing in sex chromosomes,^43^ was elevated in liver and spleen after wtITR vector transduction compared to the non-injected control mice (Fig. 4D, S3E). In contrast, the liver and spleen tissue harvested from SynITR treated mice exhibited γH2AX levels similar to the non-injected control (Fig. 4D, S3). In part, H2AX activation is typically observed in response to DNA double stranded break stress.^43,71,72^ Transient activation occurs usually within minutes to hours after the stress and γH2AX plays a role in the DNA damage repair process, possibly through retention of remodeling factors near the repair site.^72,73^ Once the DNA damage is repaired, γH2AX is dephosphorylated, usually within 6-24 hours.^74–76^ However, prolonged presence of γH2AX, termed persistent γH2AX, is associated with cell senescence,^43^ chronic inflammation associated with “senescence associated secretory phenotype proteins” including proinflammatory cytokines,^23^ and aging.^23,77,78^ To our knowledge, the data herein are the first example that wtITR rAAV infection induces persistent γH2AX in the liver and spleen of any species.

The trans-species transduction differences are another critical hurdle for AAV gene therapy. AAV serotypes or capsid mutants which transduced well in rodents often do not transduce well in the equivalent human or primate organs.^4^ Towards the development of future clinical translational applications such as AAV-scIM mediated prevention of corneal transplant rejection,^25^ the SynITR transduction was tested in a viable human corneas. The cornea is a clear tissue located on the anterior surface of the eye, and consists of the most superficial epithelium, the Bowman’s layer, the central stroma, Descemet’s membrane, and the single cell endothelium layer. The corneas can be cultured with maintaining the morphology and cell viability over 1 month,^79^ rendering it an ideal human tissue for ex vivo studies. Herein, the vectors were administered into the corneal stroma, which primarily consists of a single quiescent cell type, keratocytes, and collagen fibers.^80^ Notably, when wt and SynITR vectors were compared between two corneas from a single donor, SynITR enhanced transduction in all 4 of the individual donors, with a range of 2.6- to 16.0-fold compared to the wtITR vectors (Fig. 5). There were no obvious distinguishable differences between the individuals with bigger or moderate enhancement by SynITR in raw luciferase activity values, conditions of corneas, and demographics of the donors (including ages, races, causes of death, and past medical history) although further experimentation is necessary to confirm the lack of associations. Postmortem-injection time was slightly longer for the individuals with bigger transduction enhancement than the modest enhancement (Fig. 5); nevertheless, the underlying mechanisms with the inter-individual variation, including possibly the DNA damage response, remains to be explored. In addition, all of the corneas tested were from males and similar experiments are in progress using female tissue.

In summary, these data indicate that ITR modifications can alter the transduction efficiency consistent with several previous reports.^11,12^ Moreover, to our knowledge, the data herein are the first report to show that ITR modifications alter transduction efficiency in a tissue-specific manner following IV injection. For the SynITR, the transduction enhancement was observed in vivo, and not in various cell lines even when using the same vector preparation, highlighting that the transduction efficiency differences were not a general effect, or simply a vector titering issue. Surprisingly, the wtITR, but not SynITR, resulted in persistent γH2AX observed in the liver and spleen, which could possibly be related to epigenetic silencing^43^ of the wtITR vector genome resulting in the 7-fold higher reporter activity using SynITR vectors. Additionally, as persistent γH2AX is associated with chronic tissue inflammation, it remains possible that SynITR vectors have the capacity to decrease the liver and perhaps neurological related toxicities following wtITR administration through decreased activation of persistent γH2AX, a lower required dose due to enhanced transduction, or both. Collectively, AAV-SynITR vectors offer the opportunity for tissue-specific, more effective, low-dose, and safer AAV gene therapy. Minimally, it offers an additional context to serve as a “control” for the further elucidation of wtITR’s functions in AAV biology.

## Acknowledgement

We thank the UNC Vector Core for AAV vector production and the UNC Preclinical Research Unit for conducting the mice in vivo experiment. Appreciation is also given to Dr. Han Zongchao at UNC for giving us ARPE-19 cells and to Dr. Ula Jurkunas at The Schepens Eye Research Institute, Inc. for giving us HCEnC-21T cells. Miracles is Sight, located in Clemmons NC, is acknowledged for providing human corneas.

This study was supported in part by the Core Facility Advocacy Committee and Office of Research Technologies, UNC Chapel Hill School of Medicine (TH), and by the department of UNC Ophthalmology (MH).

## Author Contributions

T.H., N.V., E.S.N., T.K., L.S., J.J.B., and M.L.H. performed experiments and analyzed the data. M.R., R.T., P.V., and A.M.C. performed in-life animal experiments. T.H., and M.H. wrote the original draft of the manuscript and prepared the Figures. N.V., E.S.N., T.K., M.R., R.T., P.V., A.M.C., L.S., J.J.B., and R.J.S. revised the manuscript and approved the final version of the manuscript. T.H., and M.L.H. obtained the funding. M.L.H., and R.J.S. designed the concept and supervised the experiment and interpretation of the data.

