## Supplemental Materials for "Adeno-associated Virus (AAV) Synthetic Inverted Terminal Repeat Enhances Tissue Specific Transduction and Altered Vector Induced Stress Response"

**Table S1. Vector titers**

| **Vector core preparation** | **(Titer with GFP/EF1a primer/probe)** | **Full/empty (%)** | **Used experiment** |
| --- | --- | --- | --- |
| AAV2 - wtITR - GFP (prep 3) | 4.82 e^9^ vg/ul (GFP) | 94 | Verification (Fig. 1A-D),  In vitro_HEK293, ARPE-19, HCEnC-21T (Fig.2A, C, D) |
| AAV2 - SynITR - GFP (prep 3) | 1.69 e^9^ vg/ul (GFP) | 98 | (not used) |
| AAV2 - SynITR - GFP (prep 4) | 4.82 e^9^ vg/ul (GFP) | 98 | Verification (Fig. 1A-D),  In vitro_HEK293, ARPE-19, HCEnC-21T (Fig.2A, C, D) |
| AAV8 - wtITR - luciferase (prep 5) | 8.02 e^9^ vg/ul (EF1a) | 98 | Verification (Fig.1A,B, S1A)  In vivo mice (Fig. 3)  Ex vivo human cornea (Fig. 5) |
| AAV8 - SynITR - luciferase (prep 5) | 7.96 e^8^ vg/ul (EF1a) | Not determined | Verification (Fig.1A,B, S1A)  In vivo mice (Fig. 3)  Ex vivo human cornea (Fig. 5) |
| AAV8 - SynITR - luciferase (prep 6) | 1.70 e^9^ vg/ul (EF1a) | 98 | Ex vivo human cornea #4 (Fig. 5) |
| AAV9 - wtITR - GFP (prep 7) | 7.01 e^8^ vg/ul (EF1a) | 98 | (not used) |
| AAV9 - SynITR - GFP (prep 7) | 3.20 e^9^ vg/ul (EF1a) | 98 | (not used) |

A. **Vectors produced by UNC vector core**

| **In lab preparations** | **(Titer with GFP primer/probe)** | **Used experiment** |
| --- | --- | --- |
| AAV2 - wtITR - GFP(prep 1) | 8.80e^7^ vg/ul | Verification (Fig.1C),  In vitro_HepG2 (Fig.2B) |
| AAV2 - SynITR - GFP (prep 1) | 4.36e^7^ vg/ul | Verification (Fig.1C),  In vitro_HepG2 (Fig.2B) |
| AAV2 - wtITR - GFP(prep 2) | 1.00e^8^ vg/ul | Verification (Fig.1C),  In vitro_661W (Fig. 2E) |
| AAV2 - SynITR - GFP (prep 2) | 3.46e^7^ vg/ul | Verification (Fig.1C),  In vitro_661W (Fig. 2E) |

B. **Vectors produced in house by the lab**

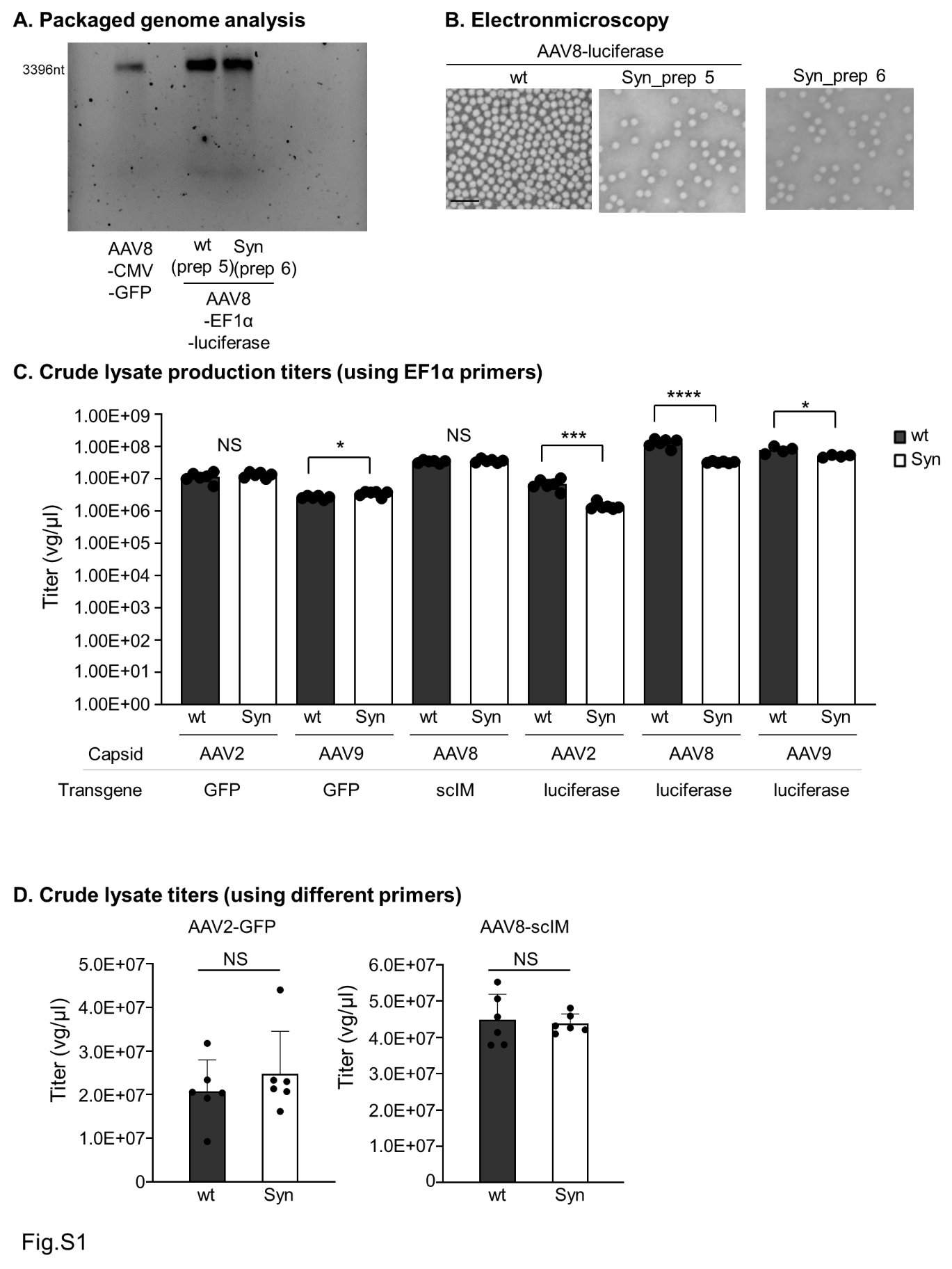

**Figure S1. Wild type or Syn ITR vector verification**. **(A)** The packaged genomes in AAV8-EF1α-wt/Syn ITR-luciferase vectors used in the human cornea ex vivo #4 (prep 5 and prep 6, Table S1) wer analyzed. Capsid packaged genomes (8.0e^9^ vg) were analyzed by alkaline gel electrophoresis and SYBR Gold staining. A wtITR vector genome (AAV8-CMV-GFP, 3,396nt) was used as a size reference. **(B)** Electron microscopy images of AAV8-luciferase wt or Syn ITR vectors (prep 5 and prep 6, Table S1). Scale bar: 100 nm. **(C, D)** Crude lysate production titers determined by ddPCR with an EF1 α primer/probe set (**C**) or with a green fluorescent protein (GFP) or single-chain immunomodulatory (sCIM) primer/probe set (**D**). *p<0.05, ***p<0.001, ****p<0.0001, NS, not statistically significant, unpaired t-test. N=6 each for AAV2-GFP, AAV9-GFP, AAV8-scIM, AAV2-luciferase and AAV8-luciferase, N=4 for AAV9-luciferase. scIM, single-chain immunomodulator.

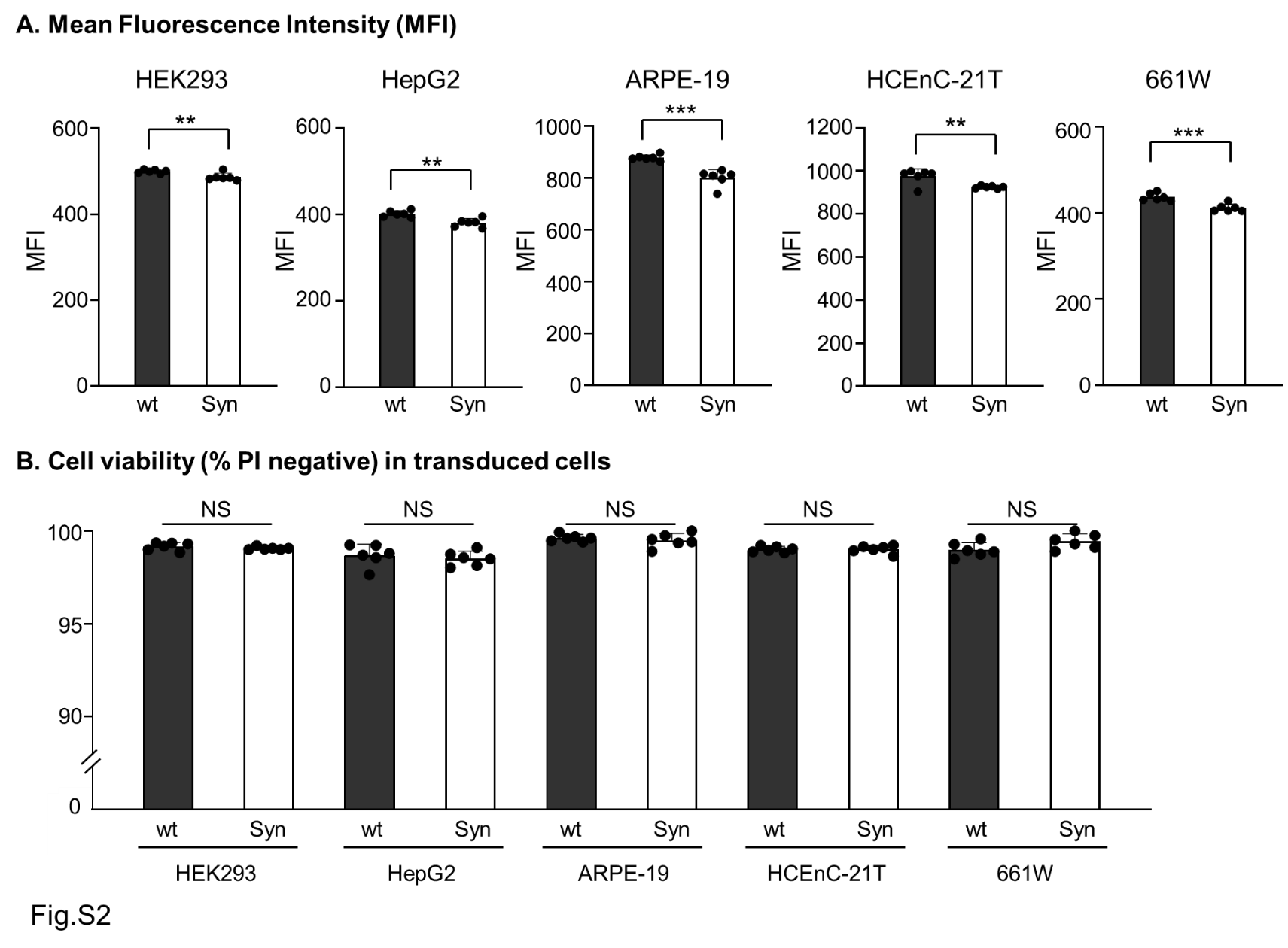
**Figure S2. Transduction efficiency of wild type or Syn ITR vectors and cell viability.** Human embryonic kidney (HEK) 293 (cells, HepG2 human hepatocellular carcinoma derived cells, adult retinal pigment epithelium (ARPE)-19 cells, highly proliferative human corneal endothelial cell line (HCEnT-21T)(**D**), or 661W mouse photoreceptor derived cells were infected with AAV2-GFP with wt or Syn ITRs at a vector genome (vg) to cell ratio of 10,000 vg/cell for HEK 293, HepG2, ARPE-19, and HCEnT-21T or 50,000 vg/cell for 661W cells. After vector addition, cells were cultured for 48hrs (HEK293) or 72hrs (HepG2, ARPE-19, HCEnT-21T, and 661W) before flow cytometry analysis. **(A)** Transduction efficiency is shown as mean intensity of transduced cells. N=6 each, **p<0.01, ***p<0.001, unpaired t-test. **(B)** Propidium iodide (PI) negative percentage in transduced cells was analyzed as a measurement of cell viability. NS, not statistically significant, unpaired t-test. N=6 each.

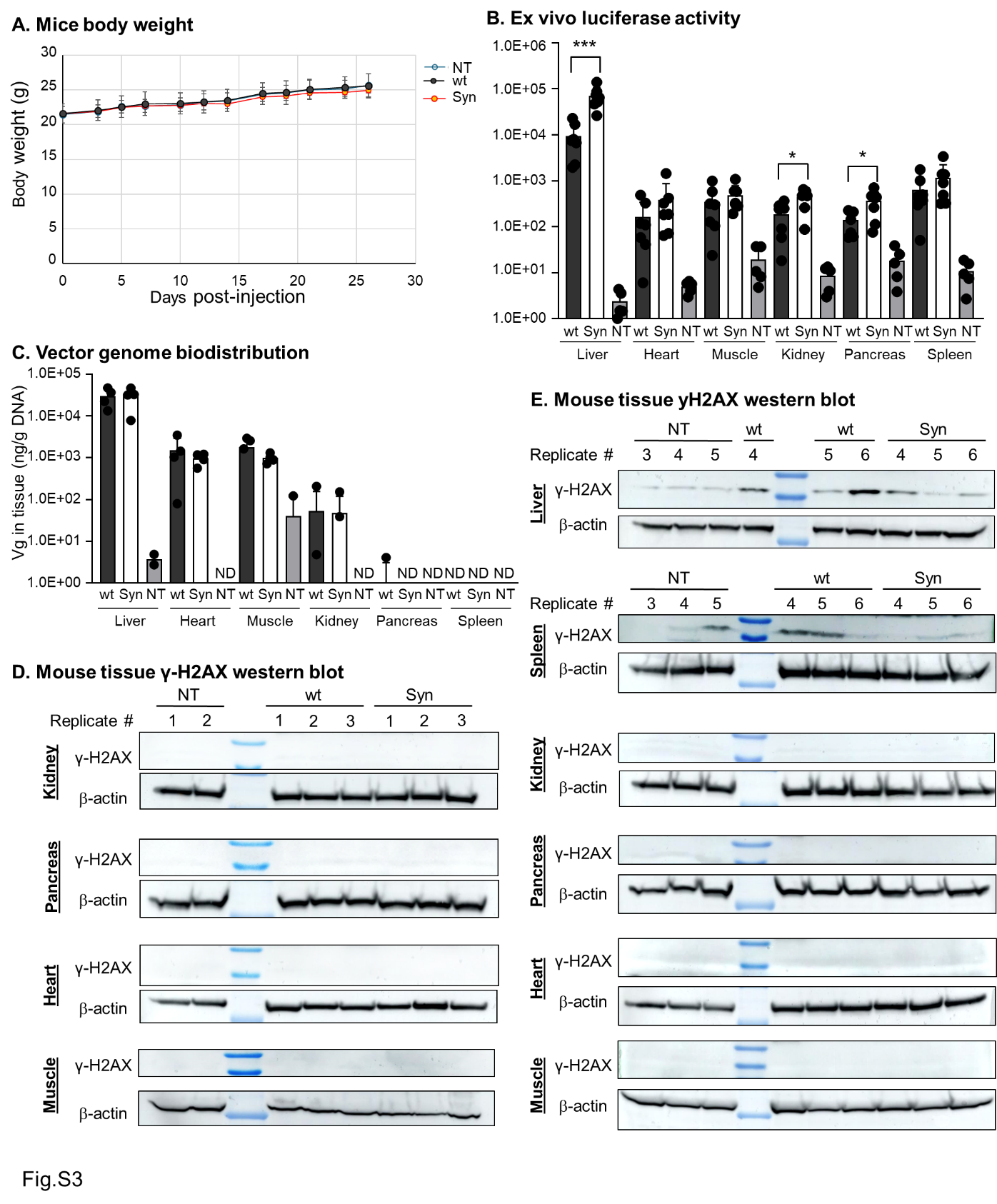

**Figure S3. AAV Syn or wt ITR vector transduction in mice.** **(A)** Body weight over time of the mice administered AAV8-EF1α-luciferase vectors with wt or Syn ITRs. NT: non-treated control mice. **(B)** Ex vivo luciferase activity in the indicated tissue. Liver, heart, muscle (tibialis anterior muscle), kidney, pancreas, and spleen were collected from mice 23 days after wt or Syn ITR vector injections. The luciferase activity was normalized to the total recovered protein concentration. N=7 for wt and Syn ITR vector injected mice, N=5 for NT. *p<0.05, ***p<0.001, Tukey’s multiple comparison test. **(C)** Vector genome biodistribution in the indicated tissue. Vector genome copy number was analyzed by quantitative polymerase chain reaction (qPCR) using primers/probe detecting the EF1α sequence. ND, not detected in all samples. N=7 for wt and Syn ITR vector injected mice, N=5 for NT. **(D, E)** Western blot analysis of phospho-histone H2AX (γ-H2AX; Ser139) of mice liver, spleen, kidney, pancreas, heart, and muscle. 50µg of protein was loaded. β-actin was used as a loading control. N=5 mice for NT, N=6 mice each for wt or Syn ITRs (Fig. **4D**, **S3D**, **S3E**). The same replicate number in each treatment group represents the sample from the same animal in Fig. 4 (**D**) andFig. S3 (**D**, **E**).

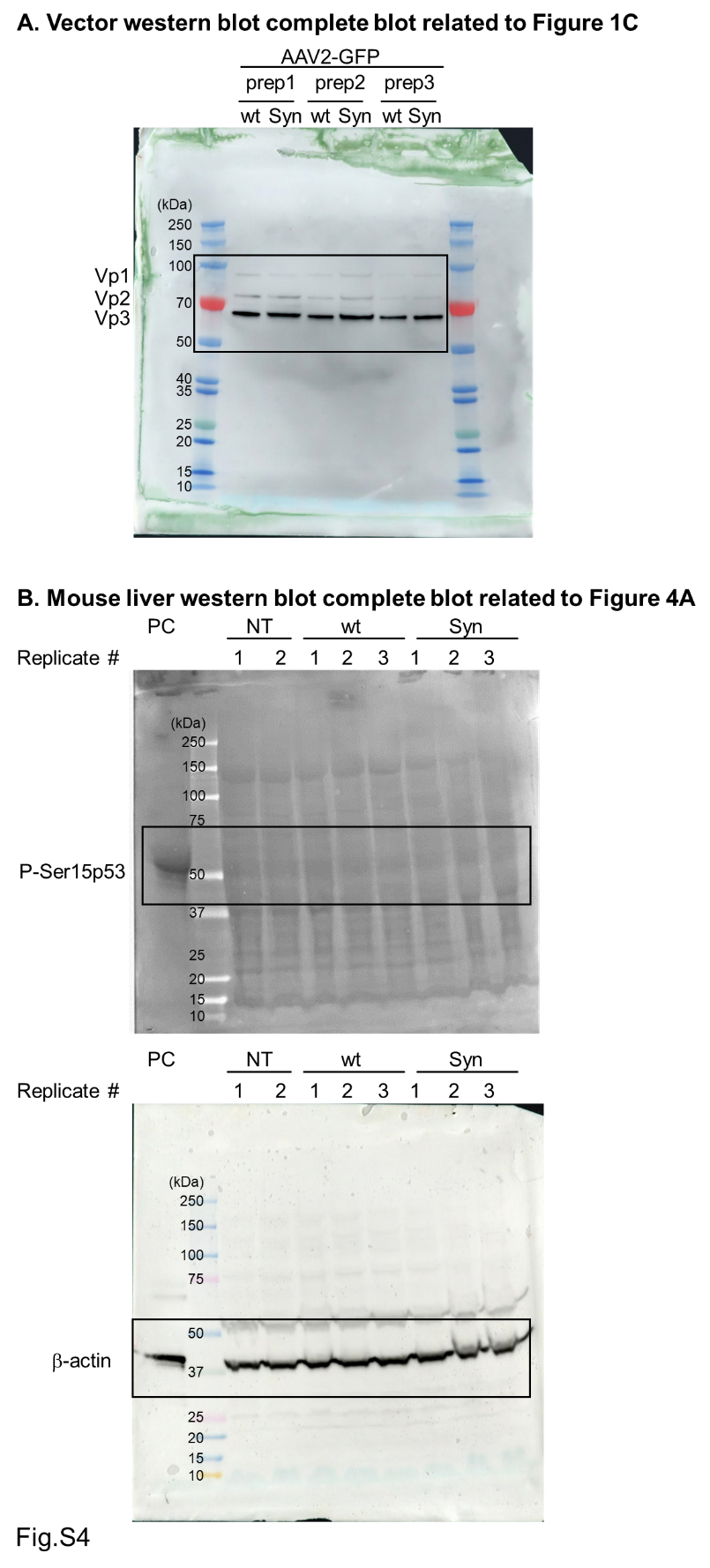

**Figure S4. Complete scans of the western blots in Figure 1C and Figure 4A. (A)** Complete scan of the western blot related to Figure 1C. AAV capsid protein western blot of vectors with wt or Syn ITRs. Same titer of the AAV2-GFP-wt or Syn ITR vectors (8.0e^8^ vg/well) were loaded and reacted with an anti-Vp1/Vp2/Vp3 antibody (capsid 2). Three independent vector productions, prepared separately by different individuals (Table S1), were tested. **(B)** Complete scan of the western blot related to Figure 4A. Wild type six week old male mice were administered with 8.0e^10^vg of AAV8-EF1α-luciferase vectors with wt or Syn ITRs by tail vein injections. Non-injected mice served as non-treated controls (NT). Liver collected from mice 23 days after the vector injections was analyzed by western blot analysis using an anti-phospho-p53 (Ser15) antibody. β-actin was used as a loading control. Etoposide treated human fibroblasts sample was used as a positive control (PC). N=2 mice for NT, N=3 mice each for wt and Syn ITR.

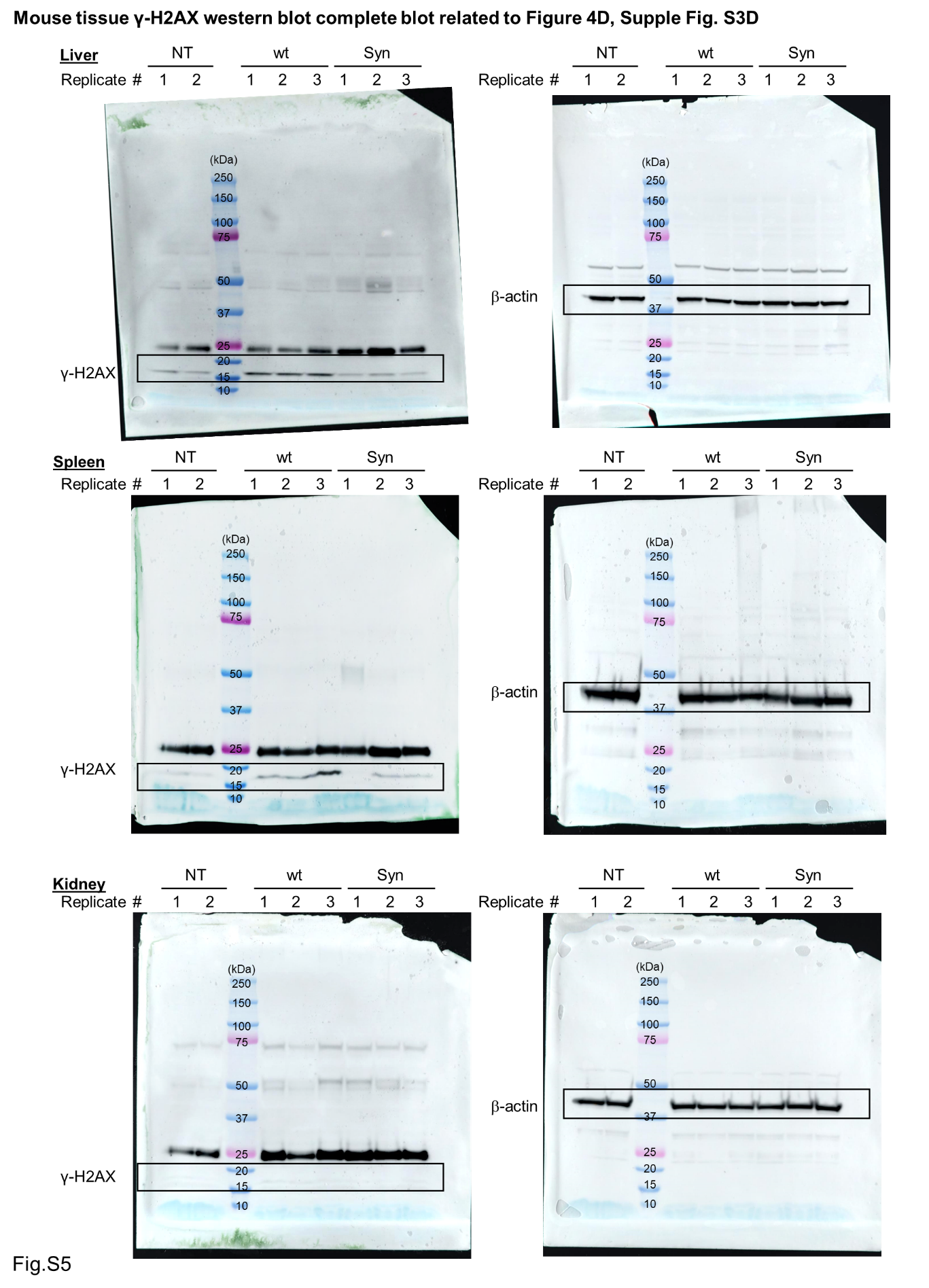

**Figure S5. Complete scans of the western blots in Figure 4D and S3D.** Wild type six week old male mice were administered with 8.0e^10^vg of AAV8-EF1α-luciferase vectors with wt or Syn ITRs by tail vein injections. Non-injected mice served as non-treated controls (NT). Liver, spleen, and kidney collected from mice 23 days after the vector injections were analyzed by western blot using an anti-phospho-histone H2AX (γ-H2AX; Ser139) antibody. β-actin was used as a loading control. N=5 mice for NT, N=6 mice each for wt and Syn ITR (Fig. **4D**, **S3**-**S8)**. The same replicate number in each treatment group represents the sample from the same animal.

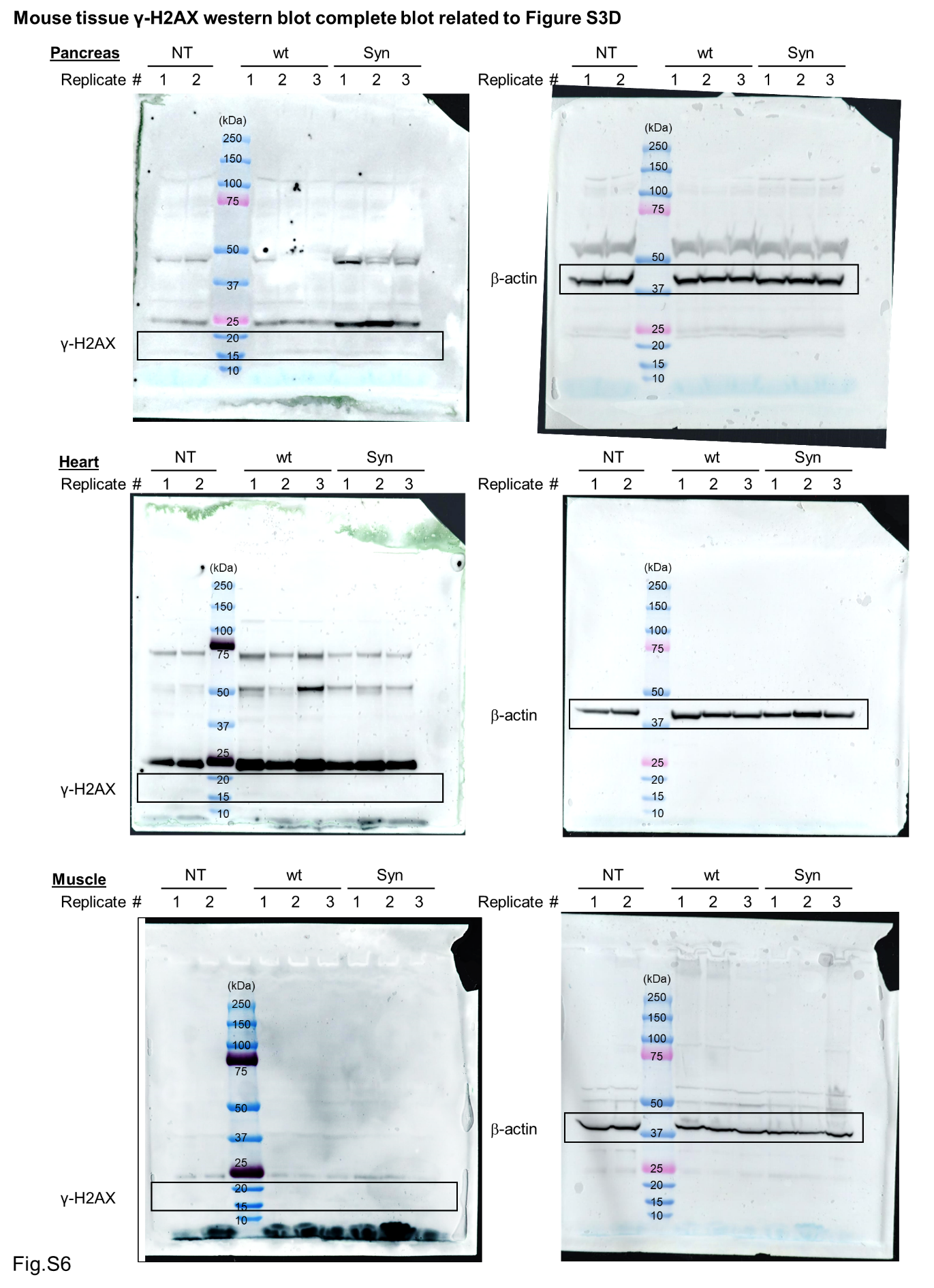

**Figure S6. Complete scans of the western blots in Figure S3D.** Wild type six week old male mice were administered with 8.0e^10^vg of AAV8-EF1α-luciferase vectors with wt or Syn ITRs by tail vein injections. Non-injected mice served as non-treated controls (NT). Pancreas, heart, and muscle (tibialis anterior muscle) collected from mice 23 days after the vector injections were analyzed by western blot with anti-phospho-histone H2AX (γ-H2AX; Ser139) antibody. β-actin was used as a loading control. N=5 mice for NT, N=6 mice each for wt and Syn ITR (Fig. **4D**, **S3**-**S8)**. The same replicate number in each treatment group represents the sample from the same animal.

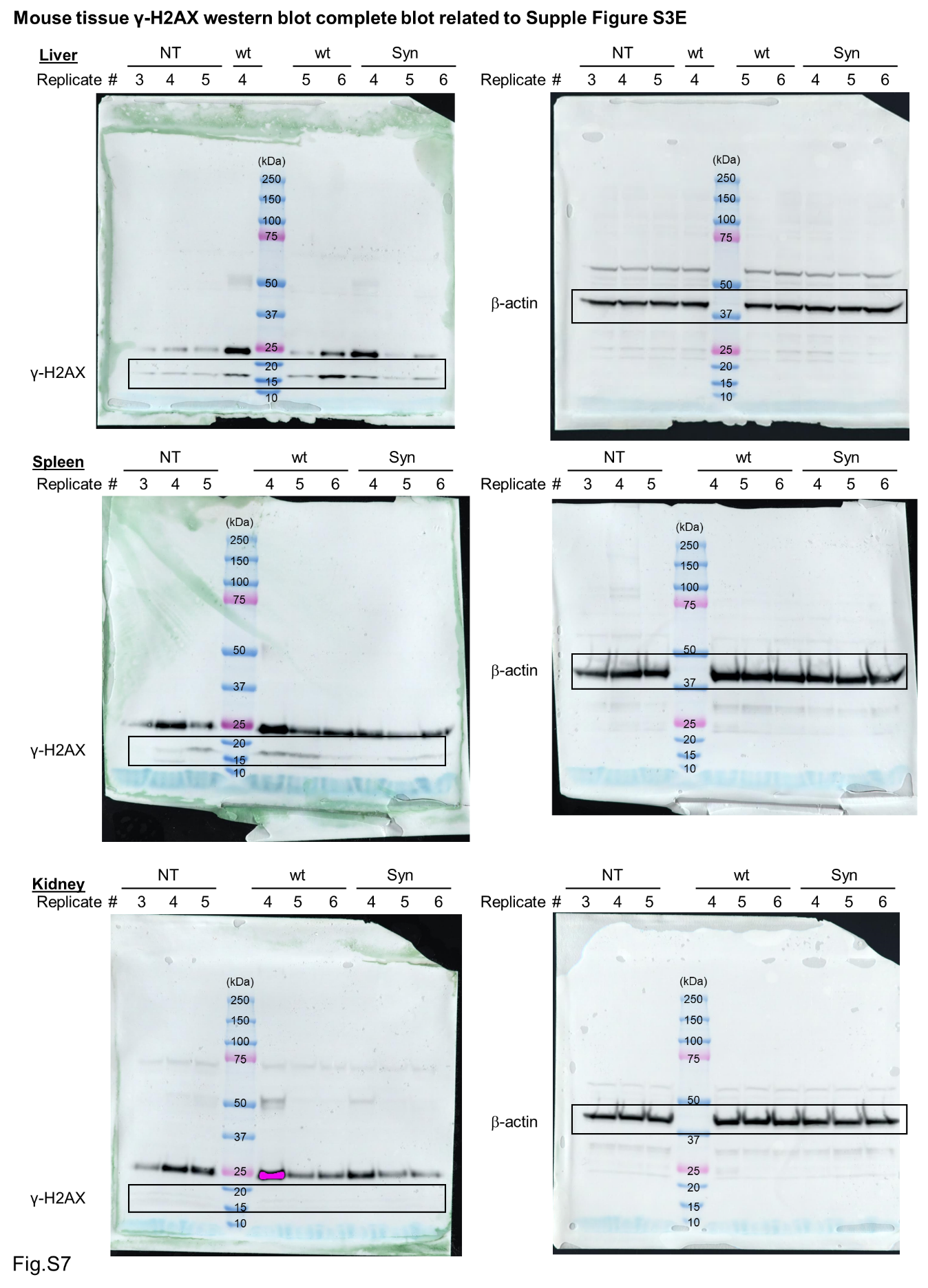

**Figure S7. Complete scans of the western blots in Figure S3E.** Wild type six week old male mice were administered with 8.0e10 vg of AAV8-EF1α-luciferase vectors with wt or Syn ITRs by tail vein injections. Non-injected mice served as non-treated controls (NT). Liver, spleen, and kidney collected from mice 23 days after the vector injections were analyzed by western blot using an anti-phospho-histone H2AX (γ-H2AX; Ser139) antibody. β-actin was used as a loading control. N=5 mice for NT, N=6 mice each for wt and Syn ITR (Fig. **4D**, **S3**-**S8)**. The same replicate number in each treatment group represents the sample from the same animal.

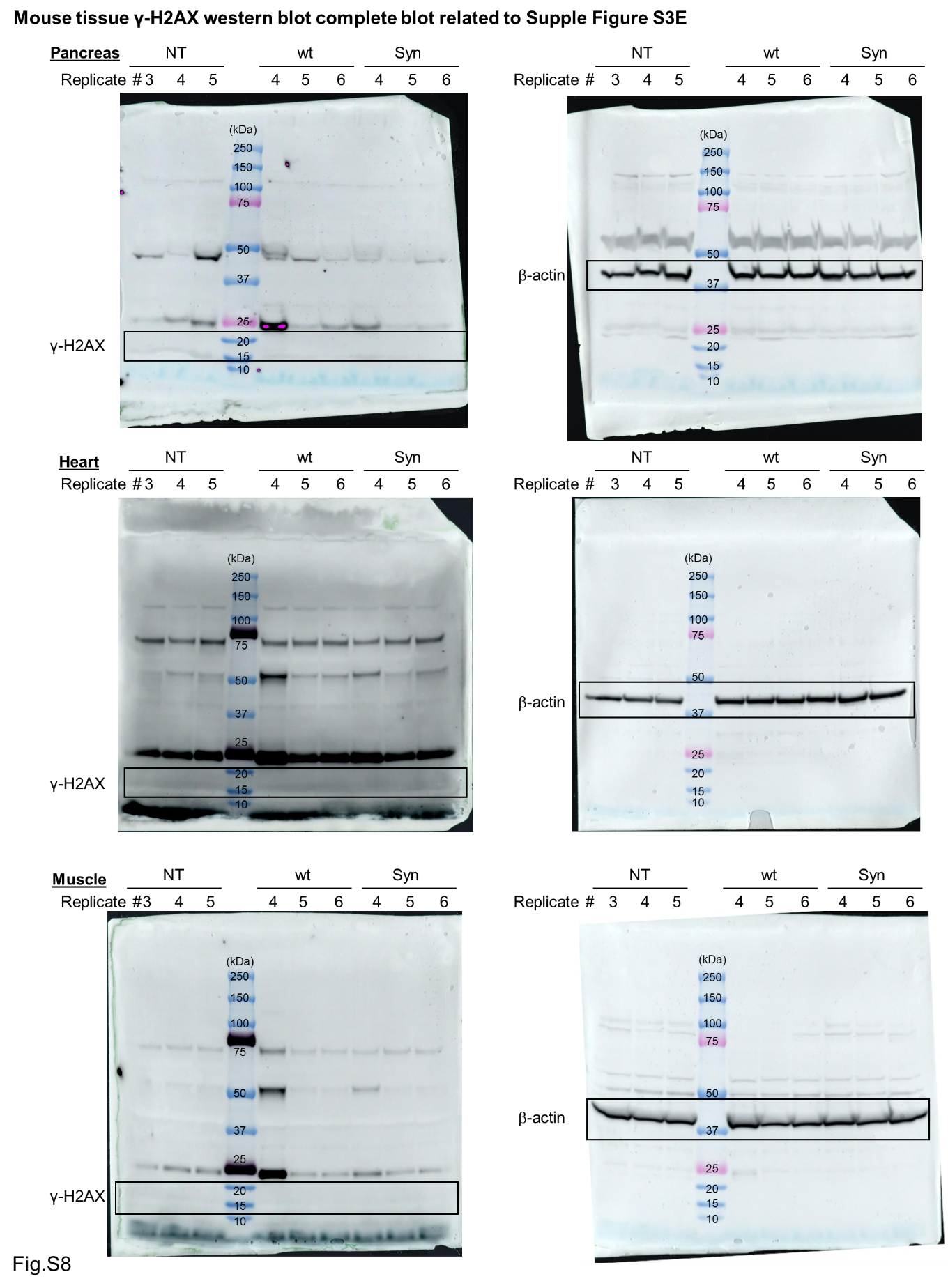

**Figure S8. Complete scans of the western blots in Figure S3E.** Wild type six week old male mice were administered with 8.0e^10^vg of AAV8-EF1α-luciferase vectors with wt or Syn ITRs by tail vein injections. Non-injected mice served as non-treated controls (NT). Pancreas, heart, and muscle (tibialis anterior muscle) collected from mice 23 days after the vector injections were analyzed by western blot using an anti-phospho-histone H2AX (γ-H2AX; Ser139) antibody. β-actin was used as a loading control. N=5 mice for NT, N=6 mice each for wt and Syn ITR (Fig. **4D**, **S3**-**S8)**. The same replicate number in each treatment group represents the sample from the same animal.
